# Mechanistic Basis for Inhibition of the Extended Spectrum Class A β-Lactamase GES-1 by Tazobactam and Enmetazobactam

**DOI:** 10.1101/2025.03.13.642993

**Authors:** Michael Beer, Philip Hinchliffe, Marko Hanževački, Christopher R. Bethel, Catherine L. Tooke, Marc W. van der Kamp, Krisztina M. Papp-Wallace, Robert A. Bonomo, Stuart Shapiro, Adrian J. Mulholland, James Spencer

## Abstract

Expression of β-lactamases is the primary form of β-lactam antibiotic resistance in Gram-negative bacteria. Enmetazobactam is a penicillanic acid sulfone (PAS) that inhibits extended spectrum β-lactamases (ESBLs) by forming an acyl-enzyme complex that eventually breaks down to an irreversible lysinoalanine crosslink. In contrast, enmetazobactam inhibits the class A carbapenemase KPC-2 *via* an acyl-enzyme that does not lead to lysinoalanine crosslink formation. This difference correlates with greater inhibitory potency of enmetazobactam against class A ESBLs, compared to carbapenemases. The GES enzymes, unlike other class A β-lactamase families, show progression from carbapenem-inhibited to carbapenem-hydrolysing phenotypes through single point mutations. We present crystal structures of GES-1, a globally disseminated ESBL, as the enmetazobactam- and tazobactam-derived acyl-enzymes. The complexes differ in the identities of their respective covalent adducts, with the catalytic Ser70 acylated by a 214 Da enmetazobactam-derived fragment, whereas tazobactam has fragmented to a 70 Da aldehyde. The tautomeric form of the enmetazobactam-derived ligand is verified by high-level QM/MM calculations, revealing the *trans*-enamine as the most thermodynamically stable tautomer, that adopts an optimised conformation that best matches the experimentally observed electron density appended to the side chain oxygen of Ser70. In contrast to previous findings for the ESBL CTX-M-15, mass spectrometry provides no evidence for lysinoalanine crosslink formation on reaction of GES-1 with (enme)tazobactam, providing further evidence that PAS inhibitors inhibit different class A β-lactamases by different mechanisms. This work reveals new details of the basis for PAS inhibition of diverse β-lactamases, and will guide development of future β-lactamase inhibitors.

## Introduction

Loss of antibiotic efficacy due to selection of resistant mutants of target bacteria is a major and growing clinical concern. Overuse of antibiotics in the clinic and in agriculture has prompted rapid evolution of resistance in many pathogenic bacteria ^1^. Emergence and dissemination of β-lactamases, that deactivate the most widely prescribed antibiotic class, β-lactams ^2^, represents a major hindrance to successful treatment of bacterial infections.

β-Lactamases are divided into four groups ^3^. Ambler classes A, C, and D comprise the serine β-lactamases (SBLs), which utilize an active site serine residue to catalyse hydrolysis of the β-lactam ring, rendering β-lactam antibiotics (and in some cases β-lactamase inhibitors containing a β-lactam moiety) inactive. Class A SBLs employ a catalytic Ser70 residue to hydrolyse β-lactam antibiotics via a covalent acyl-enzyme intermediate, with the acyl-enzyme complex hydrolysed by nucleophilic attack of a water molecule (the deacylating water), activated by Glu166, to release an inactive cleavage product ^4^. Class A enzymes have a spectrum of catalytic activity that collectively encompasses the full range of β-lactam antibiotics ^5^.

The GES (Guiana extended-spectrum) enzymes are a family of plasmid-mediated class A β-lactamases, currently comprising 59 variants ^6^, that are disseminated globally and are increasingly detected in *Pseudomonas aeruginosa, Acinetobacter baumannii* and *Klebsiella pneumoniae* ^7^. The parent enzyme, GES-1, is classified as an extended-spectrum β-lactamase (ESBL) due to an activity spectrum that encompasses penicillins and oxyimino-cephalosporins (e.g. ceftriaxone, ceftazidime, and cefepime), but not carbapenems ^8, 9^; and its susceptibility to inhibition by the mechanism-based β-lactam inhibitor clavulanic acid (IC_50_ 5-7.7 μM ^9, 10^). However, in contrast to other ESBLs, single point mutations, such as G170N (GES-2) or G170S (GES-5), endow detectable hydrolytic activity towards carbapenems, leading to carbapenem failure in patients infected by bacteria expressing GES-2/-5, or multiple other GES variants ^11–13^.

A common strategy to treat infections by bacteria expressing β-lactamases is to combine a β-lactam with a β-lactamase inhibitor (BLI) ^4, 14^. Clinically available BLIs act by forming a long-lived covalent complex with the SBL catalytic serine (Ser70 in class A SBLs), functionally inactivating the SBL and preventing efficient hydrolysis of the β-lactam antibiotic partner ^4^. Clinically utilized BLIs are subdivided into three major scaffolds: β-lactam based inhibitors (e.g. clavulanic acid, sulbactam and tazobactam) that are used most commonly; diazabicyclooctane inhibitors (e.g. avibactam); and boronate inhibitors (e.g. vaborbactam). Not unexpectedly, introduction of BLIs to the clinic has resulted in the emergence of β-lactamases that are unaffected by BLIs, whilst maintaining activity against β-lactam antibiotics ^15^.

Enmetazobactam is a β-lactam based BLI which has recently been approved for use in combination with cefepime by the European Medicines Agency (for complicated urinary tract infections (cUTI), hospital-acquired and ventilator-associated pneumonia, and bacteremic sequalae thereof) ^16^ and the U. S. Food & Drug Administration (for cUTI) ^17^. Enmetazobactam contains a penicillanic acid sulfone (PAS) scaffold ^18, 19^, like the older BLI tazobactam ^20^, but differs from the latter molecule by strategic addition of a methyl group to the triazole ring (Figure 1), rendering it zwitterionic ^20^, and accompanied by enhanced potency against bacterial cells ^19^. It is postulated that the increased potency of enmetazobactam, compared to tazobactam, derives from its ability to penetrate the bacterial outer membrane and accumulate in the periplasm of susceptible pathogens at higher concentrations per unit time ^21^.

**Figure 1:**
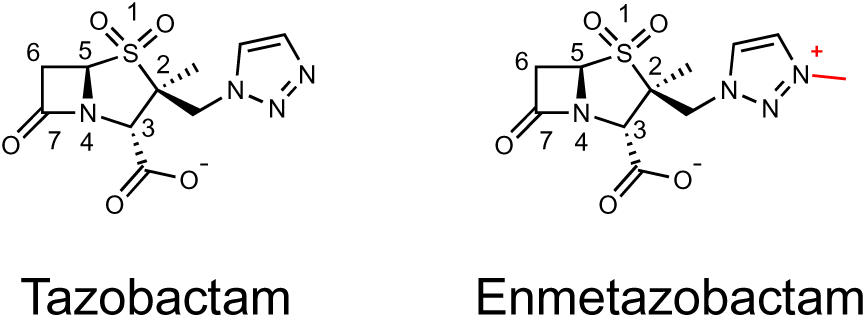
Structures of Tazobactam and Enmetazobactam. Enmetazobactam differs from tazobactam by the presence of a methyl group on the triazole ring, making it zwitterionic (red).

The reactions of PAS inhibitors with SBLs are complex, initially involving formation of an acyl-enzyme, which ramifies into multiple intermediates, as observed by crystallography, mass spectrometry and Raman spectroscopy, when tazobactam or enmetazobactam are exposed to some SBLs: e.g., class A (TEM-1, TEM-116, SHV-1, CTX-M-9, CTX-M-15, CTX-M-64, GES-2, and PC1), the class C (AmpC) enzyme CMY-2, and the class D carbapenemase OXA-48. These PAS inhibitor fragmentation pathways, after acylation and subsequent opening of the inhibitor thiazolidine ring, can result in an 88 Da hydrated aldehyde, a 70 Da aldehyde, and two separate 52 Da (vinyl ether and alkyne) adducts covalently bound to Ser70 (and to Ser130 in the case of the vinyl ether) (Figure S1) ^19, 22–24^. It is thought these different acyl-adducts may have differing reactivities with respect to deacylation ^25^. Tazobactam and enmetazobactam behave similarly on reaction with the ESBL CTX-M-15, with formation of a Ser70-Lys73 lysinoalanine crosslink as evidenced in co-crystallisation and mass spectrometry experiments (Figure S2) ^19, 26^. Mass spectrometry has shown that PAS binding to KPC-2, a class A carbapenemase, is reversible, proceeding by formation of a covalent complex upon exposure to either enmetazobactam or tazobactam, suggesting that class A carbapenemases are inhibited by enmetazobactam via covalent acyl-enzyme formation, whilst in contrast class A ESBLs are inhibited by enmetazobactam via irreversible active site crosslink formation ^19, 21–23, 27^. It is thought that differing conditions in mass spectrometry experiments favour specific fragmentation pathways, but little is known about how these fragmentation pathways may differ between proteins or whether their products are accessible by methods other than mass spectrometry ^19, 22^. Understanding the difference between ESBL and carbapenemase inhibition by BLIs is important for the development of future agents that maximise efficacy across the full range of clinically relevant enzymes.

Here, we present kinetic, mass spectrometric, and crystallographic and molecular modelling data, characterising enmetazobactam- and tazobactam-derived inhibitory complexes, that describe the reactions of the two PAS inhibitors with the class A β-lactamase GES-1. The results indicate that exposure of GES-1 crystals to enmetazobactam leads to an initial acyl-enzyme complex, most likely the *trans*-enamine breakdown intermediate (Figure S3), whereas exposure to tazobactam results in a 70 Da aldehyde acylated to Ser70, rather than a ring-opened acylated species. In contrast to earlier findings for ESBLs, formation of a lysinoalanine crosslink between Lys73 and Ser70 was not observed after soaking GES-1 with either PAS ^26^. Our results uncover mechanistic details of class A β-lactamase inhibition by enmetazobactam, which shows greater than 10-fold more potent inhibition of multiple class A ESBLs than the related tazobactam. This work may aid in guiding future development of novel PAS BLIs.

## Methods

### Enzyme Purification and Crystallisation

The GES-1 gene was synthesised with codon optimization (Eurofins, Wolverhampton, UK). The putative signal sequence of the gene was removed by amplifying from position 52 using forward primer 5’-AAGTTCTGTTTCAGGGCCCGGCGAGCGAGAAACTGACG-3’ and reverse primer of 5’-ATGGTCTAGAAAGCTTTATTTGTCGGTGGACAGGATCAG-3’ before ligation-independent insertion into the pOPIN-F T7 expression vector ^28^, digested with KpnI (NEB, Ipswich, MA, USA) and HindIII (NEB, Ipswich, MA, USA). SoluBL21 (DE3) *E. coli* (AMSBIO, San Diego, USA) cells were transformed with the resulting pOPIN-F-GES-1 plasmid and grown in 2x yeast extract tryptone (2xYT) media, supplemented with 50 µg/mL of carbenicillin. Cells were shaken at 37°C in 500 mL of liquid broth until an optical density at 600 nm (OD_600_) of 0.6-0.8 was achieved, at which time isopropyl β-**D**-1-thiogalactopyranoside (IPTG) was added to a final concentration of 0.75 mM, and the cells left shaking at 18°C overnight. Cells were harvested by centrifugation at 6,500 *x g* for 10 minutes at 4 °C and resuspended in 100 mL of buffer A (20 mM Tris pH 8.0, 300 mM NaCl) supplemented with one tablet of complete EDTA-free protease inhibitor (Roche, Basel, Switzerland), 2 µL DNase I (NEB, Ipswich, MA, USA), and 2 mg lysozyme (Sigma, Burlington, MA, USA). Cells were lysed via one passage through a cell disruptor (Constant Systems, Daventry, U.K.) at a pressure of 25 kpsi, and pelleted by centrifugation at 100,000 x *g*, at 4°C, for 1 hour. The soluble fraction was incubated, in the presence of 10 mM imidazole, with 2mL of Ni-NTA resin (Qiagen, Venlo, Netherlands) for 2 hours at 4°C. Beads were washed with 20 mL of 20 mM imidazole dissolved in buffer A, and subsequently eluted with 10 mL of 300 mM imidazole dissolved in buffer B (20 mM Tris pH 8.0, 150 mM NaCl) at 4°C. The eluent was concentrated and dialysed in an Amicon 10-kDa molecular weight (Sigma, Burlington, MA, USA) cut-off centrifugal filter, at 4°C, until the imidazole concentration reached 20 mM. Protein was then injected onto a HiLoad 16/600 Superdex 75 pg column (GE Healthcare, Amersham, UK), equilibrated with buffer B. Fractions (1 mL) were analysed using SDS-PAGE and those >95% pure were pooled and concentrated as above to 30 mg/mL.

### Crystallography, data collection, and structure determination

For ligand soaking experiments with enmetazobactam, purified GES-1 was crystallised based on the conditions reported by Smith *et al.* ^29^. Crystals were grown using sitting drop vapour diffusion in CrysChem24 plates (Hampton Research, Aliso Viejo, CA, USA) in which protein (5 mg/mL) was admixed 1:1 (4 μL final drop volume) with crystallisation mother liquor (10% (v/v) PEG 1000 and 10% (v/v) PEG 8000) and equilibrated against 500 uL mother liquor at room temperature. Crystals grew to maximum size (approx. 100 x 100 x 50 µm) within 3 weeks. Crystals were soaked in 30 mM of enmetazobactam (dissolved in mother liquor) for 1 minute or for 2.5 hours at room temperature before brief (1-5 seconds) exposure to 25 % (v/v) glycerol (4 μL drop volume) and cryo-cooling in liquid nitrogen.

For tazobactam ligand soaking, a different crystallisation condition was required due to crystals grown in the above condition dissolving upon exposure to tazobactam. Therefore, purified GES-1 (15 mg/mL) was added to a drop of 20 mM citric acid, 80 mM bis-tris propane, 10% PEG 3350 at pH 8.8 in a 1:1 ratio. Crystals were then soaked in a solution of mother liquor supplemented with 10 mM of tazobactam for 2.5 hours before brief exposure to 25% (v/v) glycerol and subsequent plunging into liquid nitrogen.

Diffraction data were collected at Diamond Light Source (Didcot, UK) on beamline I03 and indexed, integrated, scaled, and merged using Dials ^30^ in the Xia2 ^31^ processing pipeline. Phases were solved using molecular replacement with Phaser ^32^ in Phenix ^33^, with uncomplexed GES-1 (PDB ID 2QPN ^34^) as the starting structure. Structures were completed with rounds of refinement in Phenix and manual modelling in WinCoot ^35^. Ligand restraints were calculated using eLBOW in Phenix^36^. Models were validated by MolProbity ^37^ and Phenix.

### IC_50_ Determinations

IC_50_ values were obtained by following initial rates of nitrocefin (TOKU-E, Sint-Denijs-Westrem, Belgium) hydrolysis by changes in absorbance at 486 nm ^38^. The experiments were performed in kinetics buffer (10 mM HEPES, pH 7.5, 150 mM NaCl). Enmetazobactam and tazobactam (1 mM – 100 pM) were incubated with 10 nM GES-1 for 10 minutes at room temperature before addition of nitrocefin to a final concentration of 100 µM. All reactions were followed at 25°C in Greiner half-area 96-well plates in a CLARIOstar Plus microplate reader (BMG Labtech, Aylesbury, UK). IC_50_ values were calculated using GraphPad Prism 9 (GraphPad, La Jolla California, USA).

### Mass Spectrometry

Mass spectrometry data were measured as previously described ^26^ for purified GES-1 incubated for 15 minutes and for 24 h with either tazobactam or enmetazobactam. In brief, purified GES-1 was subjected to electrospray ionization-mass spectrometry (ESI-MS) after either enmetazobactam or tazobactam exposure (1:1 ratio) at *t* = 0, 15 minutes and 24 hours. All reactions were quenched with 1 % (v/v) acetonitrile and 0.1 % (v/v) formic acid in water. After quenching, samples were injected into an Acquity H class ultraperformance liquid chromatography (UPLC) instrument equipped with a 1.7 μm 2.1 mm by 100 mm Acquity UPLY ethylene bridged hybrid C_18_ column (Waters) that had been equilibrated with 0.1 % formic acid in water.

### QM/MM and QM modelling

Previous studies have shown the use of simulation to inform on crystallographic refinement^39^. To prepare systems for QM/MM geometry refinements, explicit bulk solvent was added and relaxed around the crystallographic model. In this case, all crystallographically modelled atoms (all heavy atoms from the protein, ligand and crystallographically modelled waters) were restrained whilst bulk solvent was allowed to relax around them. Chain A of the 2.5-hour enmetazobactam-soaked GES-1 crystal structure was used, keeping all crystallographic waters and one HEPES buffer molecule found near the protein-bound enmetazobactam ligand. The protonation states of all titratable residues at pH 7.4 were calculated with PROPKA 3.1 ^40^. All residues were in the standard protonation states other than Lys234 and Asp245 (in the Ambler numbering scheme) which were deprotonated and protonated, respectively. Histidine tautomers were determined by the Reduce program implemented in AmberTools21 ^41^. The all-atom AMBER ff14SB^42^ force field was used to describe protein residues. After addition of hydrogen atoms to protein-bound enmetazobactam (both the enamine and imine tautomers were modelled) and HEPES (deprotonated form), both enmetazobactam and HEPES were parameterized using General Amber Force Field (GAFF) ^43^ and their partial charges derived using the AM1-BCC method with the Antechamber package ^44^. The structure was solvated with a cubic box of TIP3P ^45^ water using the tleap module of AMBER 20 ^46^, such that the box edge was at least 10 Å from any solute atoms. No counterions were added as neutral charge of the system was not required for the subsequent QM/MM or QM geometry optimisation calculations. All heavy atoms were initially restrained to their crystallographic positions with a force constant of 500 kcal mol^-1^ Å^-2^. After an initial water energy minimization (300 steps of steepest descent with 700 steps of conjugate gradient, with all non-water heavy atoms restrained (500 kcal mol^-1^ A^-2^ restraint weight)), and a second minimization with only backbone atoms restrained (500 kcal mol^-1^ A^-2^ restraint weight), the system was subject to MD simulation for 200 ps using the NVT ensemble, with restraints maintained on all non-water heavy atoms. The system was heated to 298 K over the first 20 ps, with the temperature controlled using the Langevin thermostat with a collision frequency of 1 ps^-1^, followed by a further 5 ns of NPT simulation at constant pressure of 1.0 bar using the Berendsen barostat, maintaining the restraints on all non-water heavy atoms. All bonds involving hydrogens were constrained using the SHAKE algorithm. Long-range electrostatic interactions were treated with the Particle Mesh Ewald (PME) method with the classical non-bonded cut-off of 10 Å. A timestep of 2 fs was used in all MD simulations, which were performed using AMBER 20 software ^46^.

Final bulk-solvent equilibrated MD structures were processed to obtain non-periodic systems suitable for QM/MM calculations, used to determine the dominant tautomeric form of the acylated enmetazobactam in the enmetazobactam derived GES-1 acyl-enzyme presented here. The solvation shell around enmetazobactam was created by retaining the closest 500 water molecules to the protein-bound enmetazobactam ligand, whilst all others were removed. The active region for geometry optimisation (i.e. in which atoms were allowed to move during optimisation) was defined as all those residues whose atoms were within 6 Å of protein-bound enmetazobactam, whilst the positions of all other atoms were fixed. The QM region was defined as acylated enmetazobactam, starting from Cβ of Ser70 (Figure S4). The Cα of Ser70 was replaced by a single link hydrogen atom generated with covalent coupling. The QM region was comprised of 41 atoms, including the link hydrogen. The total charge and multiplicity of the QM region were 0 and 1, respectively. QM/MM geometry optimisation was performed with electrostatic embedding and additive scheme at the B3LYP/MM level (B3LYP/6-31G(d):AMBER ff14ffSB) including Grimme’s D3 dispersion correction and Becke-Johnson damping (D3BJ) ^47^. The QM/MM system was optimised to a minimum using the L-BFGS algorithm with a constant trust radius. Single point calculations were performed on previously optimised geometries at the B3LYP-D3(BJ)/def2-TZVP level of theory. Frequency calculation was done at the geometry optimization level of theory to confirm that the optimised structure is a true minimum. Free energy corrections at 300°K obtained from frequency calculations were included in the final electronic energies. All QM/MM calculations were performed using the ORCA 5.0.3/DL_POLY 5 interface in Py-ChemShell 21.0.3 ^48^.

Cluster models are a ‘QM-only’ methodology where only some residues or parts of some residues (e.g. the Cα atom and the side chain) are modelled, reducing the computational cost by only considering a subset of atoms involved in a protein system. The active site cluster models used here were generated using the GaussView 6.1.1 package ^49^: they included active site residue side chains (Ser70, Lys73, Asn132, Glu166, Thr237; Figure S4) and the acylated enmetazobactam in either the *trans*-enamine or imine forms. Solvent and buffer molecules were not included in the active site model (Figure S4). Geometry optimisations were performed while restraining atoms as shown in Figure S4 to maintain the positions of active site residues and acylated enmetazobactam close to the crystallographic conformation. Geometry optimisations were performed using the implicit solvation (conductor-like continuum polarisable model (CPCM), ε = 4.24) at the DFT level using the B3LYP functional and 6-31G(d) basis with Grimme’s D3 dispersion correction (important for modelling structures and interactions in proteins ^50^) and Becke-Johnson damping, set in the Gaussian 16 package ^51^. Single-point energies were calculated using the B3LYP functional and a larger (def2-TZVP) basis set including Grimme’s D3 dispersion correction and Becke-Johnson damping (Figure S4). QM calculations of isolated acylated enmetazobactam (including Ser70 side chain) also were performed at the DFT level. The B3LYP/6-31G(d) level of theory was used for geometry optimization, and the B3LYP/def2-TZVP method (both calculations included Grimme’s D3 dispersion correction and Becke-Johnson damping) for single point energy calculations.

## Results

### Enmetazobactam is a more potent inhibitor of GES-1 than tazobactam

Enmetazobactam shows strikingly different inhibitory potencies towards different class A β-lactamases, with notably lower IC_50_s for ESBLs than for carbapenemases (Table 1). With a few exceptions, reported IC_50_ values for enmetazobactam are lower than those for tazobactam for a diverse panel of class A β-lactamases. The IC_50_ of enmetazobactam for purified GES-1 is 107 nM, fourfold lower than that of tazobactam (444 nM), signalling that enmetazobactam is a stronger inhibitor of GES-1 than tazobactam. However, both tazobactam and enmetazobactam are less potent inhibitors (up to 10-fold increase in IC_50_ values) of GES-1 than of the original-spectrum β-lactamases (OSBLs) and ESBLs (Table 1) ^19, 22, 23, 52^. Indeed, the potency of tazobactam against GES-1 is similar to that for inhibitor-resistant variants of other ESBLs, e.g., TEM-30, SHV-49 and SHV-1^R244S^.

**Table 1:** IC_50_ Values of Enmetazobactam and Tazobactam against class A β-lactamases.

| Enzyme | Enzyme Classification | Enmetazobactam $IC_{50}$ (nM) | Tazobactam $IC_{50}$ (nM) | Reference |
| --- | --- | --- | --- | --- |
| GES-1 | ESBL | **107** | **444** | This Work |
| CTX-M-14 | ESBL | 6 | 100 | 19 |
| CTX-M-15 | ESBL | 7 | 6 | 19 |
| TEM-26 | ESBL | 82 | 23 | 19 |
| TEM-1 | OSBL | 7 | 30 | 19 |
| TEM-116 | OSBL | 36 | 11 | 22 |
| SHV-1 | OSBL | 8 | 26 | 19 |
| GES-2 | Carbapenemase | ND | 500 | 23 |
| SME-1 | Carbapenemase | ND | 160 | 52 |
| KPC-2 | Carbapenemase | 360 | 3700 | 19 |
| KPC-3 | Carbapenemase | 520 | 4400 | 19 |
| TEM-30 | IRT | 290 | 240 | 19 |
| SHV-1<sup>R244S</sup> | IRS | 560 | 390 | 19 |
| SHV-49 | IRS | 360 | 730 | 19 |

Key: ESBL, extended-spectrum β-lactamase; OSBL, original-spectrum β-lactamase; IRT, inhibitor-resistant TEM; IRS, inhibitor-resistant SHV; ND, not determined.

### GES-1 exposure to penicillanic acid sulfones does not lead to the formation of a lysinoalanine crosslink

To understand the modes of inhibition of GES-1 by PAS compounds, purified enzyme was exposed to tazobactam and enmetazobactam and the resultant covalent products (after 15 minutes and 24 hours incubation) analysed using mass spectrometry. Mass spectra obtained following incubation with enmetazobactam or tazobactam are very similar. At 15 minutes there is a single major peak (+52 Da), indicative of the vinyl ether product observed previously by Papp-Wallace *et al.* and by Lang *et al.* (Figure S1) ^19, 22^. At 24 hours, a single major peak at the mass of uncomplexed GES-1 indicates regeneration of the native enzyme, most likely after deacylation of the acyl-enzyme complex. Contrary to observations for CTX-M-15 (ESBL) and OXA-10 (class D oxacillinase) ^22, 26^, an 18 Da mass loss corresponding to loss of the Ser70 hydroxyl group, and formation of the Ser70-Lys73 lysinoalanine crosslink (Figure 2, Figure S2), was not observed.

**Figure 2:**
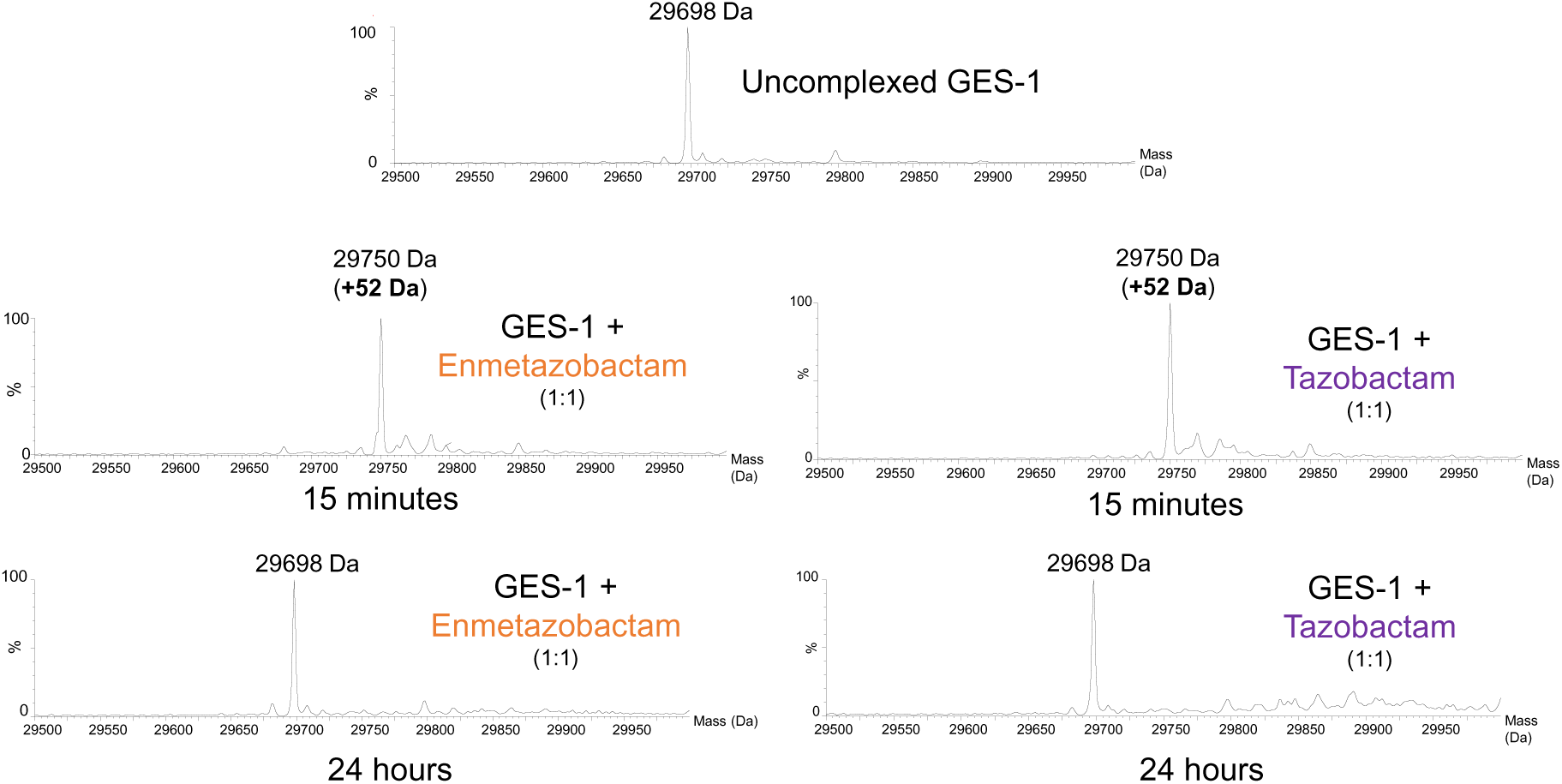
Mass spectrometric analysis of GES-1 after exposure to enmetazobactam. A) mass spectrum of the native GES-1 enzyme, in the absence of enmetazobactam exposure. B) and C) show mass spectra of GES-1 following 15-minute and 24-hour exposure to enmetazobactam and tazobactam, respectively.

### Crystallographic analysis of penicillanic acid sulfones reaction with GES-1

To visualize the products of reaction of enmetazobactam and tazobactam with purified GES-1 using X-ray crystallography, GES-1 crystals were incubated with 30 mM enmetazobactam or 10 mM tazobactam for 2.5 hours. Crystallographic data (collected at cryogenic temperature) extended to 1.23 Å and 1.36 Å resolution for the enmetazobactam and tazobactam soaks (Table S1). Both crystal forms contained two molecules in the crystallographic asymmetric unit (chains A and B, respectively), as observed in GES variants previously crystallised in the same conditions ^34, 53^. Size exclusion chromatography indicates that this is a non-physiological dimer, forming during crystallisation. We do not observe, in any of the structures presented here, electron density in the active site that could be attributed to the 52 Da vinyl ether resolved by mass spectrometry (Figure 2, S1). This matches previous reports of differences between crystallographically resolved structures and dominant adducts in solution identified by mass spectrometry ^19, 22, 26^.

Active site electron density after GES-1 exposure to tazobactam was modelled as the 70 Da aldehyde (Figure 3, S1, S5) covalently bound to Ser70. The 70 Da aldehyde covalent adduct is thought to form after fragmentation of the initial acylated inhibitor ^19, 22^. Lang *et al.* postulated that formation of the aldehyde is acid-promoted, but the conditions used here to obtain the crystal structure of the tazobactam-derived acyl-enzyme were at pH 8.8, indicating that other factors may also promote the reaction pathway for aldehyde formation. As is common for β-lactamase complexes with covalently bound β-lactam antibiotics, or classical BLIs, the C7 oxygen (Figure 1) is hydrogen-bonded by the backbone amides of Thr237 and Ser70 (Figure S6). A further hydrogen-bonding interaction occurs between the aldehyde oxygen and the amide nitrogen of Asn132 (Figure S6). No further interactions between the covalent adduct and protein were identified. Three water molecules are situated between Glu166 and the acylated aldehyde, hinting at their potential involvement in a deacylation reaction to regenerate the uncomplexed GES-1 enzyme, as observed by mass spectrometry (Figures 2, 3). A low occupancy HEPES molecule, hydrogen-bonded to Ser130, was also resolved in the active site (Figure S7). *F*_o_-*F*_c_ electron density at a packing interface could be further modelled as a molecule of intact tazobactam (Figure S9).

**Figure 3:**
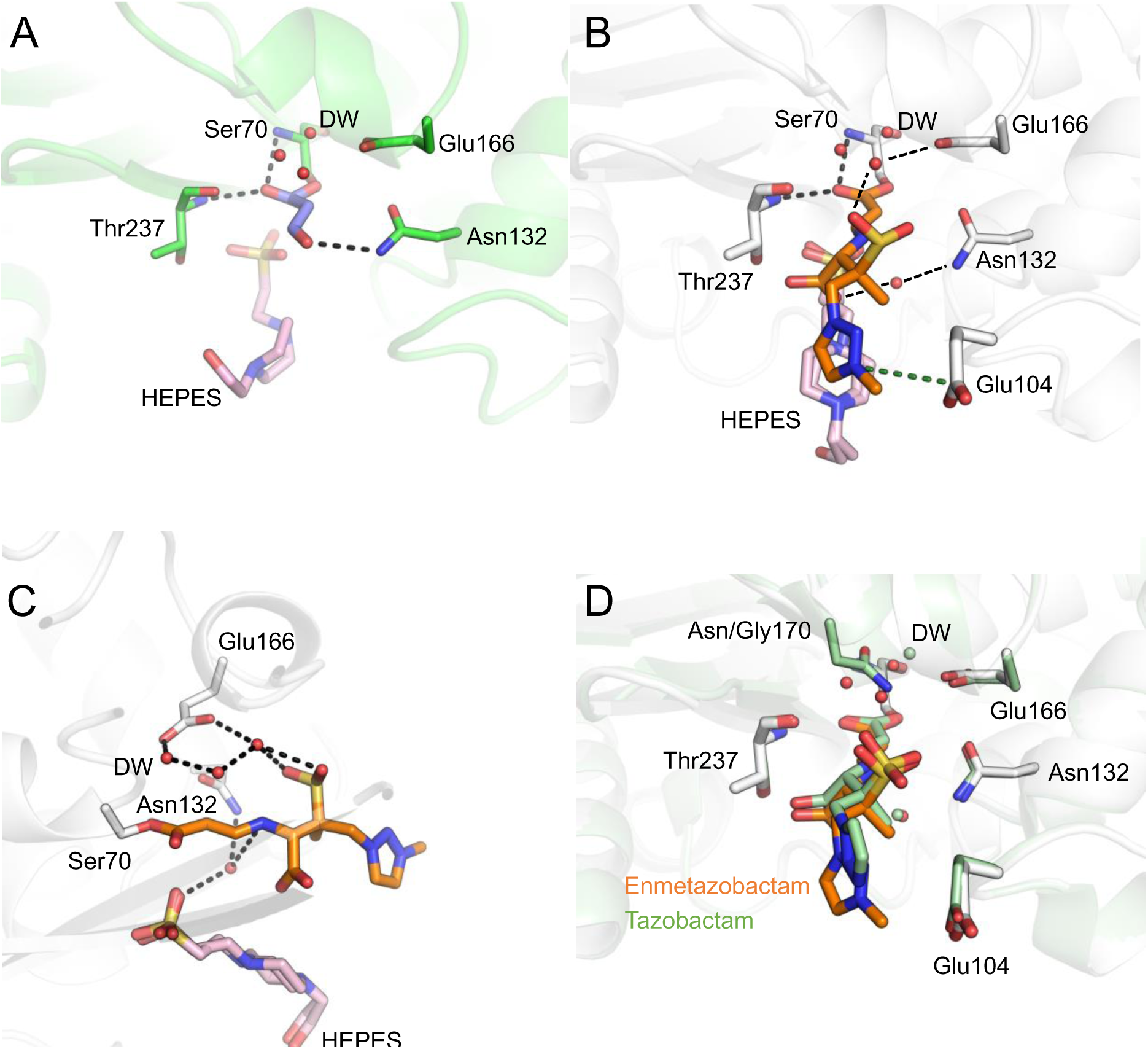
Crystallographic observation of tazobactam- and enmetazobactam-derived GES-1 acyl-enzymes. A) Tazobactam-derived GES-1 acyl-enzyme. The 70 Da covalent adduct has been modelled into the electron density (Figures S1, S5). Hydrogen bonds between the tazobactam derived covalent adduct and GES-1, including bridging interactions with water molecules, are shown (black dashes). B) Enmetazobactam-derived GES-1 acyl-enzyme. The trans-enamine tautomer was modelled into F_o_-F_c_ electron density, as supported by QM/MM structural modelling at the DFT level of theory. Hydrogen bonds between the enmetazobactam-derived covalent adduct and GES-1 are shown (black dashes) along with the electrostatic interaction between Glu104 and the positively charged methyltriazole ring (green dashes). C) A network of water molecules is resolved around the enmetazobactam derived GES-1 acyl-enzyme. Hydrogen bonds are highlighted with black dashed lines. D) Overlay of tazobactam-derived GES-2 acyl-enzyme (green protein and ligand, PDB ID 3NIA ^23^) and the enmetazobactam-derived GES-1 acyl-enzyme presented here (grey protein, orange ligand). GES-2 is a Gly170Asn mutant of GES-1, and the addition of the side chain amide in GES-2 results in sufficient electrostatic interactions to positionally stabilise a water molecule in the DW position in GES-2 crystal structures. As such, a DW is also observed in the tazobactam-derived GES-2 acyl-enzyme (PDB ID 3NIA^23^), but is positioned 1 Å away from the putative DW in the enmetazobactam GES-1 acyl-enzyme, probably as a result of steric constraints from the Gly170Asn substitution in GES-2.

X-ray diffraction data collected from GES-1 crystals after exposure to enmetazobactam revealed electron density in the GES-1 active site consistent with the imine or *trans-*enamine tautomer of the ring-opened acyl-enzyme product (Figures S1, S3, S9). Modelling of the *trans*-enamine tautomer followed the observation that the linear difference electron density more closely matched a planar *trans*-enamine conformation than an imine. The acyl-enzyme product is thought to form early in the reaction pathway of PAS compounds with class A SBLs and has been documented in structures of SBL complexes with PAS inhibitor-derived species ^19, 20, 22, 23, 26^. No electron density consistent with formation of the 52 Da vinyl ether, detected by mass spectrometry (Figure 2, S2) and expected to be bound to both Ser70 and Ser130, was observed. Thus, the pathway for vinyl ether formation may be promoted by the conditions of mass spectrometry experiments, a conclusion reached previously about other breakdown fragments ^22^; while, in the case of enmetazobactam, the acylated sulphone may be stabilized by conditions within the crystal.

The enmetazobactam-derived acyl-enzyme forms only a few electrostatic interactions with active site residues of GES-1 (Figure S6). The C7 oxygen (Figure 1) sits within the oxyanion hole (formed by the backbone amides of Ser70 and Thr237, Figure 3), consistent with structures of other β-lactam antibiotics and BLIs covalently bound to class A β-lactamases ^4^. No other enzyme:enmetazobactam hydrogen bonds are formed in the acyl-enzyme complex. Similarly, few enzyme:PAS inhibitor hydrogen bonds were observed in the crystal structure of the GES-1^G170N^ variant, GES-2, in complex with tazobactam derived inhibitory species ^23^. However, a number of water molecules in the GES-1:enmetazobactam complex structures presented here appear to make bridging interactions connecting the acylated enmetazobactam moiety and enzyme active site residues (Figure 3B, C), including between the enmetazobactam sulfone and the side chain oxygen of Glu166, and with the backbone carbonyl of Gly170. Furthermore, the positive charge around the triazole ring (conferred by the additional methyl group of enmetazobactam compared to tazobactam; Figure 1) forms a weak electrostatic interaction with the negatively charged side chain of Glu104 (distances between the positively charged nitrogen of the enmetazobactam triazole ring and Cδ of Glu104 4.3 Å and 3.8 Å, in chains A and B, respectively, Figure 3).

An additional noteworthy observation in the crystal structures presented here, compared to other GES-1 structures, is the presence of a water molecule in the deacylating position ^4^ (Figure 3). Such a water molecule has not been observed previously in crystal structures of GES-1, probably due to the inability of Gly170 (a position commonly occupied by Asn in other class A β-lactamases, including those listed in Table 1) to participate in hydrogen bonds ^11, 34^. This water molecule is surrounded by a further network of water molecules, also not observed in other GES-1 structures, with a bridging water between the sulphone group of the enmetazobactam-derived acyl-adduct and the putative deacylating water. The observation of a putative deacylating water is consistent with mass spectrometry results that show GES-1 as capable of turning over the species formed on reaction with enmetazobactam. The putative deacylating water is in an optimal position for proton transfer to Glu166 (or possibly Lys73) and to make a nucleophilic attack upon the C7 carbon (Figure 1) of Ser70-bound species in the first step of the deacylation reaction.

In both the tazobactam- and enmetazobactam-derived complexes, further electron density in the active site could be modelled as a molecule of HEPES (present during GES-1 purification). In the enmetazobactam-derived complex, this HEPES molecule was modelled in dual conformation (Figure S7). Electron density in the same position was also observed in the presented structure of otherwise uncomplexed GES-1, supporting the decision to model this as HEPES (Figure S7, Table S1). This density is in a similar position to the non-covalent product modelled in the previously reported GES-2:tazobactam complex structure ^23^, but superposition of the respective electron density maps and structures does not suggest that HEPES would be a better fit in the GES-2:tazobactam complex structure, or that the previously described non-covalent product would be a better fit to the current data (Figure S8).

### QM/MM calculations indicate that the *trans-*enamine is the dominant tautomer in GES-1:enmetazobactam complexes

Due to resolution limits and the inherent complications of modelling mixed populations of tautomers, the presented, and previously deposited, crystal structures of class A β-lactamase:(enme)tazobactam complexes are, alone, insufficiently accurate to determine whether the favoured acyl-enzyme tautomeric state is the *trans*-enamine or the imine^26^. Molecular dynamics simulations of CTX-M-15:tazobactam complexes, and previous analyses of dihedral angles of SBL:tazobactam complexes ^19^, support the imine as the favoured form. However, other crystallographic and spectroscopic studies suggest that, whilst there may be a mixed population of both tautomers, the *trans*-enamine is the dominant species ^23, 54^. Here, modelling either the imine or *trans*-enamine species into experimental electron density gave similar real-space correlation coefficient (RSCC) values: 0.94/0.95 (chains A/B) for the *trans*-enamine, and 0.94/0.94 for the imine, respectively, and therefore do not distinguish between them. To resolve this uncertainty, QM/MM calculations (using the ORCA 5.0.3:DL_POLY 5 interface in Py-ChemShell, see Methods) were employed to investigate the structural features and energetic properties of the two possible acyl-enzyme tautomeric states (Figure S1, S3). Visual inspection of QM/MM optimised geometries indicates that the *trans*-enamine tautomer of enmetazobactam more closely resembles the crystallographically modelled enmetazobactam-derived acyl-enzyme than does the imine (Figure S12). Detailed analysis of the enmetazobactam C2-C3-C4 and C3-C4-N5 angles, C2-C3-C4-N5 torsion angle, and distances between enmetazobactam N5-HEPES O1, and between enmetazobactam N5 – enmetazobactam O10, confirms that the QM/MM optimised *trans*-enamine, rather than the QM/MM optimised imine, more closely matches the models refined into the electron density (Figure S13, Table S2). Further, QM calculations on active site models (Figure S4) of acylated enmetazobactam, and QM calculations of isolated acylated enmetazobactam, indicated that the *trans*-enamine is the lowest energy tautomer (Figure S3, Table S3). Taken together, these data suggest that the crystal structure of the GES-1:enmetazobactam complex predominantly contains the *trans*-enamine. Subsequently, the crystal structure of the enmetazobactam-derived acyl-enzyme was modelled using a ring-opened enmetazobactam-derived covalent adduct, attached to Ser70, in the *trans*-enamine tautomer (Figure 3, S2, S3).

## Discussion

GES-1 is an ESBL capable of hydrolysing a range of penicillin and cephalosporin substrates, but not carbapenems. However, GES-1 is unusual among ESBLs in that it contains active site features, such as a disulphide bridge between Cys69 and Cys238, which are present in class A carbapenemases such as KPC-2 and SFC-1, and that are thought to be determinants of carbapenemase activity ^55^. Despite this, GES-1 lacks carbapenemase activity. Nevertheless, GES-1 possesses a mixture of residues that promote or inhibit carbapenemase activity at positions proposed to configure the active site electric field in carbapenem-derived acyl-enzyme complexes ^56^. Notably and unusually, some single point mutations (G170S or G170N) convert GES-1 into a carbapenemase ^11, 12, 57^.

PAS inhibitors, in particular enmetazobactam, inhibit ESBLs with greater potency than class A carbapenemases (Table 1). This is probably related to different inhibition mechanisms: enmetazobactam inhibits carbapenemases via formation of a stable acyl-enzyme complex that, however, turns over within 24 hours, whilst ESBLs are inhibited by eventual irreversible active site crosslink formation ^26^. Here, we provide mass spectrometric and crystallographic evidence that enmetazobactam is a moderately potent inhibitor of the ESBL GES-1 that acts through formation of the type of stable acyl-enzyme observed previously in class A carbapenemases, rather than through the irreversible active site crosslink formation identified previously as the mechanism of enmetazobactam inhibition of other ESBLs.

Enmetazobactam is a more potent inhibitor of GES-1 than tazobactam *in vitro* (Table 1), despite the structural similarity of these two PAS inhibitors. The four-fold lower inhibition potency of tazobactam, compared to enmetazobactam, suggests that cationicity localized to the triazole ring of enmetazobactam has either a significant effect on the active site chemistry of GES-1 or changes the strength of interactions between the enzyme and inhibitor. The greater potency of enmetazobactam towards GES-1, compared to tazobactam, contrasts with data for other β-lactamases (Table 1): TEM-26 and TEM-116 are more strongly inhibited by tazobactam, whilst both PAS inhibitors are approximately equipotent against CTX-M-15 and TEM-30. Furthermore, IC_50_ values for the inhibitors towards GES-1 presented here are closer to those reported previously for inhibitor-resistant TEM and SHV variants than for other class A ESBLs, indicating that GES enzymes may have intrinsically reduced susceptibility to mechanism-based inhibitors, including PAS compounds and clavulanic acid (IC_50_ values for GES-1 inhibition by clavulanic acid range between 5 - 7.7 μM ^8–10^).

Although the enmetazobactam-derived acyl-enzyme makes few direct interactions with residues in the GES-1 active site, bridging interactions are evident with several bound water molecules. The presence of multiple water molecules in the active site appear to form a hydrogen bonding network between the catalytic Glu166 and acyl group of the covalent bound adduct. This network, where one water molecule also forms a hydrogen bond to the sulphone group of the enmetazobactam derived acyl-adduct may also form electrostatic interactions as part of a network of water molecules between the acyl group of the covalent adduct and the catalytic Glu166, resulting in a water molecule in position for a deacylation reaction, required to complete enmetazobactam turnover (Figure 3); regeneration of the uncomplexed enzyme is observed by mass spectrometry after 24-hour incubation of the complex (Figure 2). The interaction between Glu104 and the positively charged methyltriazole ring of the enmetazobactam-derived acyl-enzyme also may contribute to the enhanced potency of enmetazobactam, compared to tazobactam. This favourable electrostatic interaction, not possible with the uncharged triazole ring of tazobactam, may increase the stability of the enmetazobactam acyl-enzyme, preventing fragmentation (which is seen for the tazobactam-derived acyl-enzyme (Figure 3)), and resulting in more potent inhibition, as reflected in the IC_50_ values. However, the presence of a deacylating water, well positioned for nucleophilic attack on the enmetazobactam-derived acyl-enzyme, may indicate why GES-1 is inherently less susceptible to inhibition by enmetazobactam than other class A ESBLs (Table 1). The presence of a HEPES molecule in the active site of the crystallised complex may provide additional electrostatic stabilization of acylated enmetazobactam (Figure 3). The impact of this HEPES molecule in stabilizing a particular covalent adduct, or of promoting specific breakdown pathways *in crystallo*, is unknown.

Our QM/MM, active site QM and acyl-adduct only QM models (Figure S4), using DFT calculations, showed that the *trans*-enamine is the dominant tautomeric form of acylated enmetazobactam. The magnitude of the stability difference between the *trans*-enamine and the imine depends on the hydrogen bonding of two nearby crystallographic water molecules, according to the QM/MM calculations (Figure S14). The proximity of these two water molecules to the enmetazobactam N5 nitrogen (Figure S13) involved in tautomerization, and their impact on the relative energies of the two tautomers (Table S3), suggest that they may be important for, and perhaps participate actively in, tautomerization. The impact of the inclusion of two water molecules, that may not have been included in small active site models (Figure S4), demonstrates the advantage of using a full enzyme QM/MM model with explicit solvent, such as that offered by the ORCA 5.3.0:DL_POLY 5 interface in Py-ChemShell, which includes the protein and solvent environment, for modelling protein-ligand systems,. Active site only or single molecule QM calculations may not contain all of the important interactions that are important in an enzyme-ligand system. This work also further indicates the usefulness of employing QM/MM geometry optimization as part of crystallographic refinement in cases where ligand electron density is inconclusive ^58^.

In conclusion, our work here presents an understanding of how enmetazobactam and tazobactam inhibit the clinically concerning GES-1 β-lactamase and extends evidence that enmetazobactam inhibits different class A β-lactamases in mechanistically different ways. We show that lysinoalanine cross-link formation is not the sole mechanism of inhibition for class A ESBLs ^26^ by (enme)tazobactam and, subsequently, that reversible covalent inhibition by these compounds is not specific to class A carbapenemases. We observe that the newly approved enmetazobactam can inhibit the clinically emerging β-lactamase, GES-1, by formation of a 214 Da inhibitory acyl-enzyme. This inhibition mechanism differs from our observation of the inhibition of GES-1 by tazobactam, which forms a 70 Da inhibitory complex after fragmentation of the initial bound adduct, as observed crystallographically. QM calculations indicate that the population of enmetazobactam-derived 214 Da covalent adducts acylated in the GES-1 active site is dominated by the *trans*-enamine, rather than the imine, tautomer. The results of this study contribute to growing evidence that GES β-lactamases behave distinctly from other class A β-lactamases, having features characteristic of both ESBL and carbapenemase enzymes and that extend beyond the ability of class members to progress to carbapenemase activity through single amino acid substitutions on the parent enzyme ^11, 12, 56, 59^. There may then be potential for differences between existing, or newly emerging, GES variants in enmetazobactam susceptibility. To our knowledge, this is as yet little explored for GES-producing clinical isolates, although a recent study that included GES-1, GES-2 and GES-5 identified no variant dependence of cefepime/enmetazobactam susceptibility for recombinant *E. coli* or *Pseudomonas aeruginosa* ^60^. The greater concentration of enmetazobactam than tazobactam in the bacterial periplasm may reduce the impact on bacterial susceptibility of variant-specific differences in inhibitory potency. These considerations notwithstanding, the work presented here evidences that enmetazobactam inhibits class A β-lactamases by multiple mechanisms and suggests that strategies designed to promote fragmentation to more stable covalent adducts or evade β-lactamase-catalysed turnover, by e.g. disrupting interactions involving the deacylating water molecule, may further enhance potency towards targets such as the GES enzymes.

## Author Information

### Author Contributions

MB, PH, JS, RAB and AJM conceived the experiments. JS and AJM supervised the PhD work of MB. MB performed all wet-lab experiments. PH and CLT supervised MB in protein purification, crystallization and IC_50_ determinations. CRB and KMPW performed the mass spectrometry experiments. MH and MWVdK advised on simulations and assisted MB in QM/MM geometry optimization and single point energy calculations. MB, PH and MH contributed to data interpretation. MB wrote the initial manuscript draft, with contributions from SS and revisions and approval by all authors.

### Present Addresses

KMPW – JMI Laboratories, a subsidiary of Element Materials Technology, North Liberty, Iowa, USA

### Data Availability

Input files and ligand parameters for all QM/MM geometry optimisations and single point energy calculations are made available at the University of Bristol Research Data Repository (https://data.bris.ac.uk/). All crystal structures presented here have atomic coordinates and structure factors deposited to the Worldwide Protein Data Bank (PDB; wwwpdb.org) under accession codes 9ENV, 9ENW, 9ENX and 9ENY.

## Acknowledgements

The authors would like to thank Diamond Light Source (beamline I03, proposals 23269 and 31440) and the beamline scientists for their assistance and service that enabled the X-ray diffraction data presented here to be collected. Allecra Therapeutics provided enmetazobactam samples used in this work. The authors also would like to thank members of the EPSRC project FEHybCat for useful discussions surrounding QM/MM geometry optimisations and single point energy calculations in Py-ChemShell.

## Current Addresses

KMPW – JMI Laboratories/Element-Iowa City, North Liberty, Iowa, USA

## References

(1) Mulchandani, R.; Wang, Y.; Gilbert, M.; Van Boeckel, T. P. Global trends in antimicrobial use in food-producing animals: 2020 to 2030. PLOS Global Public Health 2023, 3 (2), e0001305. DOI: 10.1371/journal.pgph.0001305. Shallcross, L. J.; Davies, D. S. Antibiotic overuse: a key driver of antimicrobial resistance. Br J Gen Pract 2014, 64 (629), 604–605. DOI: 10.3399/bjgp14X682561 From NLM.

(2) Bush, K. Past and Present Perspectives on β-Lactamases. Antimicrob Agents Chemother 2018, 62(10). DOI: 10.1128/aac.01076-18 From NLM. Bush, K.; Bradford, P. A. β-Lactams and β-Lactamase Inhibitors: An Overview. Cold Spring Harb Perspect Med 2016, 6 (8). DOI: 10.1101/cshperspect.a025247 From NLM.

(3) Ambler, R. Structure of Beta-Lactamases. Philosophical Transactions of the Royal Society of London. Series B, Biological Sciences 1980, 289 (1036), 321–331.

(4) Tooke, C. L.; Hinchliffe, P.; Bragginton, E. C.; Colenso, C. K.; Hirvonen, V. H. A.; Takebayashi, Y.; Spencer, J. β-Lactamases and β-Lactamase Inhibitors in the 21st Century. J Mol Biol 2019, 431 (18), 3472–3500. DOI: 10.1016/j.jmb.2019.04.002 From NLM.

(5) Hao, G. F.; Yang, G. F.; Zhan, C. G. Structure-based methods for predicting target mutation-induced drug resistance and rational drug design to overcome the problem. Drug Discov Today 2012, 17 (19-20), 1121–1126. DOI: 10.1016/j.drudis.2012.06.018 From NLM.

(6) Naas, T.; Oueslati, S.; Bonnin, R. A.; Dabos, M. L.; Zavala, A.; Dortet, L.; Retailleau, P.; Iorga, B. I. Beta-lactamase database (BLDB) - structure and function. J Enzyme Inhib Med Chem 2017, 32 (1), 917–919. DOI: 10.1080/14756366.2017.1344235 PubMed.

(7) Weldhagen Gerhard, F.; Poirel, L.; Nordmann, P. Ambler Class A Extended-Spectrum β-Lactamases in Pseudomonas aeruginosa: Novel Developments and Clinical Impact. Antimicrobial Agents and Chemotherapy 2003, 47(8), 2385–2392. DOI: 10.1128/aac.47.8.2385-2392.2003 (acccessed 2024/03/08). Literacka, E.; Izdebski, R.; Urbanowicz, P.; Żabicka, D.; Klepacka, J.; Sowa-Sierant, I.; Żak, I.; Garus-Jakubowska, A.; Hryniewicz, W.; Gniadkowski, M. Spread of Klebsiella pneumoniae ST45 Producing GES-5 Carbapenemase or GES-1 Extended-Spectrum β-Lactamase in Newborns and Infants. Antimicrob Agents Chemother 2020, 64 (9). DOI: 10.1128/aac.00595-20 From NLM. Bonnin Rémy, A.; Jousset Agnès, B.; Urvoy, N.; Gauthier, L.; Tlili, L.; Creton, E.; Cotellon, G.; Arthur, F.; Dortet, L.; Naas, T. Detection of GES-5 Carbapenemase in Klebsiella pneumoniae, a Newcomer in France. Antimicrobial Agents and Chemotherapy 2017, 61 (3), 10.1128/aac.02263-02216. DOI: 10.1128/aac.02263-16 (acccessed 2024/03/08). Bogaerts, P.; Naas, T.; El Garch, F.; Cuzon, G.; Deplano, A.; Delaire, T.; Huang, T. D.; Lissoir, B.; Nordmann, P.; Glupczynski, Y. GES extended-spectrum β-lactamases in Acinetobacter baumannii isolates in Belgium. Antimicrob Agents Chemother 2010, 54 (11), 4872–4878. DOI: 10.1128/aac.00871-10 From NLM. Bonnin Rémy, A.; Rotimi Vincent, O.; Al Hubail, M.; Gasiorowski, E.; Al Sweih, N.; Nordmann, P.; Poirel, L. Wide Dissemination of GES-Type Carbapenemases in Acinetobacter baumannii Isolates in Kuwait. Antimicrobial Agents and Chemotherapy 2013, 57 (1), 183–188. DOI: 10.1128/aac.01384-12 (acccessed 2024/03/08).

(8) Lee, S. H.; Jeong, S. H. Nomenclature of GES-type extended-spectrum beta-lactamases. In Antimicrob Agents Chemother, Vol. 49; 2005; pp 2148; author reply 2148-2150.

(9) Poirel, L.; Le Thomas, I.; Naas, T.; Karim, A.; Nordmann, P. Biochemical sequence analyses of GES-1, a novel class A extended-spectrum beta-lactamase, and the class 1 integron In52 from Klebsiella pneumoniae. Antimicrob Agents Chemother 2000, 44 (3), 622–632. DOI: 10.1128/aac.44.3.622-632.2000 From NLM.

(10) Bontron, S.; Poirel, L.; Nordmann, P. In vitro prediction of the evolution of GES-1 β-lactamase hydrolytic activity. Antimicrob Agents Chemother 2015, 59 (3), 1664–1670. DOI: 10.1128/aac.04450-14 From NLM.

(11) Smith, C. A.; Frase, H.; Toth, M.; Kumarasiri, M.; Wiafe, K.; Munoz, J.; Mobashery, S.; Vakulenko, S. B. Structural basis for progression toward the carbapenemase activity in the GES family of β-lactamases. J Am Chem Soc 2012, 134 (48), 19512–19515. DOI: 10.1021/ja308197j From NLM.

(12) Stewart, N. K.; Smith, C. A.; Frase, H.; Black, D. J.; Vakulenko, S. B. Kinetic and Structural Requirements for Carbapenemase Activity in GES-type beta-lactamases. Biochemistry 2015, 54 (2), 588–597. DOI: 10.1021/bi501052t.

(13) Poirel, L.; Weldhagen, G. F.; Naas, T.; De Champs, C.; Dove, M. G.; Nordmann, P. GES-2, a class A beta-lactamase from Pseudomonas aeruginosa with increased hydrolysis of imipenem. Antimicrob Agents Chemother 2001, 45 (9), 2598–2603. DOI: 10.1128/aac.45.9.2598-2603.2001 From NLM. Bae, I. K.; Lee, Y. N.; Jeong, S. H.; Hong, S. G.; Lee, J. H.; Lee, S. H.; Kim, H. J.; Youn, H. Genetic and biochemical characterization of GES-5, an extended-spectrum class A beta-lactamase from Klebsiella pneumoniae. Diagn Microbiol Infect Dis 2007, 58 (4), 465–468. DOI: 10.1016/j.diagmicrobio.2007.02.013 From NLM.

(14) Weldhagen, G. F. Eiamphungporn, W.; Schaduangrat, N.; Malik, A. A.; Nantasenamat, C. Tackling the Antibiotic Resistance Caused by Class A β-Lactamases through the Use of β-Lactamase Inhibitory Protein. Int J Mol Sci 2018, 19 (8), 2222. DOI: 10.3390/ijms19082222 PubMed.

(15) Chaïbi, E. B.; Sirot, D.; Paul, G.; Labia, R. Inhibitor-resistant TEM β-lactamases: phenotypic, genetic and biochemical characteristics. Journal of Antimicrobial Chemotherapy 1999, 43 (4), 447–458. DOI: 10.1093/jac/43.4.447 (acccessed 9/18/2024).

(16) Agency, E. M. https://www.ema.europa.eu/en/documents/smop-initial/chmp-summary-positive-opinion-exblifep_en.pdf (accessed.

(17) Administration, U. S. F. D. https://www.accessdata.fda.gov/drugsatfda_docs/appletter/2024/216165Orig1s000ltr.pdf (accessed.

(18) Arer, V.; Kar, D. Biochemical exploration of β-lactamase inhibitors. Frontiers in Genetics 2023, 13, Review. DOI: 10.3389/fgene.2022.1060736.

(19) Papp-Wallace, K. M.; Bethel, C. R.; Caillon, J.; Barnes, M. D.; Potel, G.; Bajaksouzian, S.; Rutter, J. D.; Reghal, A.; Shapiro, S.; Taracila, M. A.;, et al. Beyond Piperacillin-Tazobactam: Cefepime and AAI101 as a Potent β-Lactam-β-Lactamase Inhibitor Combination. Antimicrob Agents Chemother 2019, 63 (5). DOI: 10.1128/aac.00105-19 From NLM.

(20) Papp-Wallace, K. M.; Bethel, C. R.; Barnes, M. D.; Rutter, J. D.; Taracila, M. A.; Bajaksouzian, S.; Jacobs, M. R.; Bonomo, R. A. AAI101, a Novel β-Lactamase Inhibitor: Microbiological and Enzymatic Profiling. In Open Forum Infect Dis, Vol. 4; © The Author 2017. Published by Oxford University Press on behalf of Infectious Diseases Society of America., 2017; p S375.

(21) Papp-Wallace, K. M.; Bethel, C. R.; Caillon, J.; Barnes, M. D.; Potel, G.; Bajaksouzian, S.; Rutter, J. D.; Reghal, A.; Shapiro, S.; Taracila, M. A.;, et al. Beyond Piperacillin-Tazobactam: Cefepime and AAI101 as a Potent β-Lactam−β-Lactamase Inhibitor Combination. Antimicrobial Agents and Chemotherapy 2019, 63 (5), 10.1128/aac.00105-00119. DOI: doi:10.1128/aac.00105-19.

(22) Lang, P. A.; Raj, R.; Tumber, A.; Lohans, C. T.; Rabe, P.; Robinson, C. V.; Brem, J.; Schofield, C. J. Studies on enmetazobactam clarify mechanisms of widely used β-lactamase inhibitors. Proceedings of the National Academy of Sciences 2022, 119 (18), e2117310119. DOI: doi:10.1073/pnas.2117310119.

(23) Frase, H.; Smith, C. A.; Toth, M.; Champion, M. M.; Mobashery, S.; Vakulenko, S. B. Identification of products of inhibition of GES-2 beta-lactamase by tazobactam by x-ray crystallography and spectrometry. J Biol Chem 2011, 286 (16), 14396–14409. DOI: 10.1074/jbc.M110.208744 From NLM.

(24) Bonomo, R. A.; Liu, J.; Chen, Y.; Ng, L.; Hujer, A. M.; Anderson, V. E. Inactivation of CMY-2 β-lactamase by tazobactam: initial mass spectroscopic characterization. Biochimica et Biophysica Acta (BBA) - Protein Structure and Molecular Enzymology 2001, 1547 (2), 196–205. DOI: 10.1016/S0167-4838(01)00175-3. Pagan-Rodriguez, D.; Zhou, X.; Simmons, R.; Bethel, C. R.; Hujer, A. M.; Helfand, M. S.; Jin, Z.; Guo, B.; Anderson, V. E.; Ng, L. M.;, et al. Tazobactam Inactivation of SHV-1 and the Inhibitor-resistant Ser130 → Gly SHV-1 β-Lactamase: INSIGHTS INTO THE MECHANISM OF INHIBITION*. Journal of Biological Chemistry 2004, 279 (19), 19494–19501. DOI: 10.1074/jbc.M311669200. Bethel Christopher, R.; Taracila, M.; Shyr, T.; Thomson Jodi, M.; Distler Anne, M.; Hujer Kristine, M.; Hujer Andrea, M.; Endimiani, A.; Papp-Wallace, K.; Bonnet, R.;, et al. Exploring the Inhibition of CTX-M-9 by β-Lactamase Inhibitors and Carbapenems. Antimicrobial Agents and Chemotherapy 2011, 55 (7), 3465–3475. DOI: 10.1128/aac.00089-11 (acccessed 2024/02/27). Thomson, J. M.; Distler, A. M.; Bonomo, R. A. Overcoming Resistance to β-Lactamase Inhibitors: Comparing Sulbactam to Novel Inhibitors against Clavulanate Resistant SHV Enzymes with Substitutions at Ambler Position 244. Biochemistry 2007, 46 (40), 11361–11368. DOI: 10.1021/bi700792a. Cheng, Q.; Xu, C.; Chai, J.; Zhang, R.; Wai chi Chan, E.; Chen, S. Structural Insight into the Mechanism of Inhibitor Resistance in CTX-M-199, a CTX-M-64 Variant Carrying the S130T Substitution. ACS Infectious Diseases 2020, 6 (4), 577–587. DOI: 10.1021/acsinfecdis.9b00345.

(25) Fritz, R. A.; Alzate-Morales, J. H.; Spencer, J.; Mulholland, A. J.; van der Kamp, M. W. Multiscale Simulations of Clavulanate Inhibition Identify the Reactive Complex in Class A β-Lactamases and Predict the Efficiency of Inhibition. Biochemistry 2018, 57 (26), 3560–3563. DOI: 10.1021/acs.biochem.8b00480 From NLM.

(26) Hinchliffe, P.; Tooke, C. L.; Bethel, C. R.; Wang, B.; Arthur, C.; Heesom, K. J.; Shapiro, S.; Schlatzer, D. M.; Papp-Wallace, K. M.; Bonomo, R. A.;, et al. Penicillanic Acid Sulfones Inactivate the Extended-Spectrum β-Lactamase CTX-M-15 through Formation of a Serine-Lysine Cross-Link: an Alternative Mechanism of β-Lactamase Inhibition. mBio 2022, 13 (3), e0179321. DOI: 10.1128/mbio.01793-21 From NLM.

(27) Kalp, M.; Totir, M. A.; Buynak, J. D.; Carey, P. R. Different intermediate populations formed by tazobactam, sulbactam, and clavulanate reacting with SHV-1 beta-lactamases: Raman crystallographic evidence. J Am Chem Soc 2009, 131 (6), 2338–2347. DOI: 10.1021/ja808311s From NLM.

(28) Berrow, N. S.; Alderton, D.; Sainsbury, S.; Nettleship, J.; Assenberg, R.; Rahman, N.; Stuart, D. I.; Owens, R. J. A versatile ligation-independent cloning method suitable for high-throughput expression screening applications. Nucleic Acids Res 2007, 35 (6), e45. DOI: 10.1093/nar/gkm047 From NLM.

(29) Smith, C. A.; Frase, H.; Toth, M.; Kumarasiri, M.; Wiafe, K.; Munoz, J.; Mobashery, S.; Vakulenko, S. B.

(30) Winter, G.; Waterman, D. G.; Parkhurst, J. M.; Brewster, A. S.; Gildea, R. J.; Gerstel, M.; Fuentes-Montero, L.; Vollmar, M.; Michels-Clark, T.; Young, I. D.;, et al. DIALS: implementation and evaluation of a new integration package. Acta Crystallogr D Struct Biol 2018, 74 (Pt 2), 85–97. DOI: 10.1107/S2059798317017235 PubMed.

(31) Winter, G. xia2: an expert system for macromolecular crystallography data reduction. Journal of Applied Crystallography 2010, 43 (1), 186–190. DOI: doi:10.1107/S0021889809045701.

(32) McCoy, A. J.; Grosse-Kunstleve, R. W.; Adams, P. D.; Winn, M. D.; Storoni, L. C.; Read, R. J. Phaser crystallographic software. Journal of Applied Crystallography 2007, 40 (4), 658–674. DOI: doi:10.1107/S0021889807021206.

(33) Adams, P. D.; Afonine, P. V.; Bunkóczi, G.; Chen, V. B.; Davis, I. W.; Echols, N.; Headd, J. J.; Hung, L.-W.; Kapral, G. J.; Grosse-Kunstleve, R. W.;, et al. PHENIX: a comprehensive Python-based system for macromolecular structure solution. Acta crystallographica. Section D, Biological crystallography 2010, 66 (Pt 2), 213–221. DOI: 10.1107/S0907444909052925 PubMed.

(34) Smith, C. A.; Caccamo, M.; Kantardjieff, K. A.; Vakulenko, S. Structure of GES-1 at atomic resolution: insights into the evolution of carbapenamase activity in the class A extended-spectrum [beta]-lactamases. Acta Crystallographica Section D 2007, 63 (9), 982–992. DOI: doi:10.1107/S0907444907036955.

(35) Emsley, P.; Lohkamp, B.; Scott, W. G.; Cowtan, K. Features and development of Coot. Acta Crystallogr D Biol Crystallogr 2010, 66 (Pt 4), 486–501. DOI: 10.1107/s0907444910007493 From NLM.

(36) Moriarty, N. W.; Grosse-Kunstleve, R. W.; Adams, P. D. electronic Ligand Builder and Optimization Workbench (eLBOW): a tool for ligand coordinate and restraint generation. Acta Crystallogr D Biol Crystallogr 2009, 65 (Pt 10), 1074–1080. DOI: 10.1107/s0907444909029436 From NLM.

(37) Williams, C. J.; Headd, J. J.; Moriarty, N. W.; Prisant, M. G.; Videau, L. L.; Deis, L. N.; Verma, V.; Keedy, D. A.; Hintze, B. J.; Chen, V. B.;, et al. MolProbity: More and better reference data for improved all-atom structure validation. Protein Sci 2018, 27 (1), 293–315. DOI: 10.1002/pro.3330 From NLM.

(38) O’Callaghan, C. H.; Morris, A.; Kirby, S. M.; Shingler, A. H. Novel method for detection of beta-lactamases by using a chromogenic cephalosporin substrate. Antimicrob Agents Chemother 1972, 1 (4), 283–288. DOI: 10.1128/aac.1.4.283 From NLM.

(39) Fonseca, F.; Chudyk, E. I.; van der Kamp, M. W.; Correia, A.; Mulholland, A. J.; Spencer, J. The Basis for Carbapenem Hydrolysis by Class A β-Lactamases: A Combined Investigation using Crystallography and Simulations. Journal of the American Chemical Society 2012, 134 (44), 18275–18285. DOI: 10.1021/ja304460j.

(40) Søndergaard, C. R.; Olsson, M. H. M.; Rostkowski, M.; Jensen, J. H. Improved Treatment of Ligands and Coupling Effects in Empirical Calculation and Rationalization of pKa Values. Journal of Chemical Theory and Computation 2011, 7 (7), 2284–2295. DOI: 10.1021/ct200133y.

(41) Case, D. A.; Aktulga, H. M.; Belfon, K.; Ben-Shalom, I.; Brozell, S. R.; Cerutti, D. S.; Cheatham III, T. E.; Cruzeiro, V. W. D.; Darden, T. A.; Duke, R. E. Amber 2021; University of California, San Francisco, 2021.

(42) Maier, J. A.; Martinez, C.; Kasavajhala, K.; Wickstrom, L.; Hauser, K. E.; Simmerling, C. ff14SB: Improving the Accuracy of Protein Side Chain and Backbone Parameters from ff99SB. Journal of Chemical Theory and Computation 2015, 11 (8), 3696–3713. DOI: 10.1021/acs.jctc.5b00255.

(43) Wang, J.; Wolf, R. M.; Caldwell, J. W.; Kollman, P. A.; Case, D. A. Development and testing of a general amber force field. J Comput Chem 2004, 25 (9), 1157–1174. DOI: 10.1002/jcc.20035 From NLM.

(44) Wang, J.; Wang, W.; Kollman, P. A.; Case, D. A. Automatic atom type and bond type perception in molecular mechanical calculations. Journal of Molecular Graphics and Modelling 2006, 25 (2), 247–260. DOI: 10.1016/j.jmgm.2005.12.005.

(45) Jorgensen, W. L.; Chandrasekhar, J.; Madura, J. D.; Impey, R. W.; Klein, M. L. Comparison of simple potential functions for simulating liquid water. The Journal of Chemical Physics 1983, 79 (2), 926–935. DOI: 10.1063/1.445869 (acccessed 1/29/2024).

(46) AMBER 2020; University of California: San Francisco, 2020. (accessed.

(47) Grimme, S.; Ehrlich, S.; Goerigk, L. Effect of the damping function in dispersion corrected density functional theory. Journal of Computational Chemistry 2011, 32 (7), 1456–1465. DOI: 10.1002/jcc.21759 (acccessed 2024/02/12). Grimme, S.; Hansen, A.; Brandenburg, J. G.; Bannwarth, C. Dispersion-Corrected Mean-Field Electronic Structure Methods. Chemical Reviews 2016, 116 (9), 5105–5154. DOI: 10.1021/acs.chemrev.5b00533.

(48) Lu, Y.; Sen, K.; Yong, C.; Gunn, D. S. D.; Purton, J. A.; Guan, J.; Desmoutier, A.; Abdul Nasir, J.; Zhang, X.; Zhu, L.;, et al. Multiscale QM/MM modelling of catalytic systems with ChemShell. Physical Chemistry Chemical Physics 2023, 25 (33), 21816–21835, 10.1039/D3CP00648D. DOI: 10.1039/D3CP00648D. Lu, Y.; Farrow, M. R.; Fayon, P.; Logsdail, A. J.; Sokol, A. A.; Catlow, C. R. A.; Sherwood, P.; Keal, T. W. Open-Source, Python-Based Redevelopment of the ChemShell Multiscale QM/MM Environment. Journal of Chemical Theory and Computation 2019, 15 (2), 1317–1328. DOI: 10.1021/acs.jctc.8b01036.

(49) GaussView; Semichem Inc: Shawnee Mission, KS, 2016. (accessed.

(50) Lonsdale, R.; Harvey, J. N.; Mulholland, A. J. Inclusion of Dispersion Effects Significantly Improves Accuracy of Calculated Reaction Barriers for Cytochrome P450 Catalyzed Reactions. The Journal of Physical Chemistry Letters 2010, 1 (21), 3232–3237. DOI: 10.1021/jz101279n.

(51) Gaussian 16 Rev. C.01; Wallingford, CT, 2016. (accessed.

(52) Bush, K.; Macalintal, C.; Rasmussen, B. A.; Lee, V. J.; Yang, Y. Kinetic interactions of tazobactam with beta-lactamases from all major structural classes. Antimicrob Agents Chemother 1993, 37 (4), 851–858. DOI: 10.1128/aac.37.4.851 From NLM.

(53) Bebrone, C.; Bogaerts, P.; Delbrück, H.; Bennink, S.; Kupper Michaël, B.; Rezende de Castro, R.; Glupczynski, Y.; Hoffmann Kurt, M. GES-18, a New Carbapenem-Hydrolyzing GES-Type β-Lactamase from Pseudomonas aeruginosa That Contains Ile80 and Ser170 Residues. Antimicrobial Agents and Chemotherapy 2013, 57 (1), 396–401. DOI: 10.1128/aac.01784-12 (acccessed 2024/07/29).

(54) Padayatti, P. S.; Helfand, M. S.; Totir, M. A.; Carey, M. P.; Hujer, A. M.; Carey, P. R.; Bonomo, R. A.; van den Akker, F. Tazobactam Forms a Stoichiometric trans-Enamine Intermediate in the E166A Variant of SHV-1 β-Lactamase: 1.63 Å Crystal Structure. Biochemistry 2004, 43 (4), 843–848. DOI: 10.1021/bi035985m. Pagan-Rodriguez, D.; Zhou, X.; Simmons, R.; Bethel, C. R.; Hujer, A. M.; Helfand, M. S.; Jin, Z.; Guo, B.; Anderson, V. E.; Ng, L. M.;, et al. Tazobactam Inactivation of SHV-1 and the Inhibitor-resistant Ser^130^ &#x2192; Gly SHV-1 &#x3b2;-Lactamase: INSIGHTS INTO THE MECHANISM OF INHIBITION *. Journal of Biological Chemistry 2004, 279 (19), 19494–19501. DOI: 10.1074/jbc.M311669200 (acccessed 2024/01/03). Grigorenko, V. G.; Petrova, T. E.; Carolan, C.; Rubtsova, M. Y.; Uporov, I. V.; Pereira, J.; Chojnowski, G.; Samygina, V. R.; Lamzin, V. S.; Egorov, A. M. Crystal structures of the molecular class A [beta]-lactamase TEM-171 and its complexes with tazobactam. Acta Crystallographica Section D 2022, 78 (7), 825–834. DOI: doi:10.1107/S2059798322004879. Kuzin, A. P.; Nukaga, M.; Nukaga, Y.; Hujer, A.; Bonomo, R. A.; Knox, J. R. Inhibition of the SHV-1 β-Lactamase by Sulfones: Crystallographic Observation of Two Reaction Intermediates with Tazobactam. Biochemistry 2001, 40 (6), 1861–1866. DOI: 10.1021/bi0022745. Sun, T.; Bethel, C. R.; Bonomo, R. A.; Knox, J. R. Inhibitor-Resistant Class A β-Lactamases: Consequences of the Ser130-to-Gly Mutation Seen in Apo and Tazobactam Structures of the SHV-1 Variant. Biochemistry 2004, 43 (44), 14111–14117. DOI: 10.1021/bi0487903. Tassoni, R.; Blok, A.; Pannu, N. S.; Ubbink, M. New Conformations of Acylation Adducts of Inhibitors of β-Lactamase from Mycobacterium tuberculosis. Biochemistry 2019, 58 (7), 997–1009. DOI: 10.1021/acs.biochem.8b01085. Totir, M. A.; Padayatti, P. S.; Helfand, M. S.; Carey, M. P.; Bonomo, R. A.; Carey, P. R.; van den Akker, F. Effect of the Inhibitor-Resistant M69V Substitution on the Structures and Populations of trans-Enamine β-Lactamase Intermediates. Biochemistry 2006, 45 (39), 11895–11904. DOI: 10.1021/bi060990m.

(55) Chudyk, E. I.; Beer, M.; Limb, M. A. L.; Jones, C. A.; Spencer, J.; van der Kamp, M. W.; Mulholland, A. J. QM/MM Simulations Reveal the Determinants of Carbapenemase Activity in Class A β-Lactamases. ACS Infect Dis 2022, 8 (8), 1521–1532. DOI: 10.1021/acsinfecdis.2c00152 From NLM.

(56) Jabeen, H.; Beer, M.; Spencer, J.; van der Kamp, M. W.; Bunzel, H. A.; Mulholland, A. J. Electric Fields Are a Key Determinant of Carbapenemase Activity in Class A β-Lactamases. ACS Catalysis 2024, 7166–7172. DOI: 10.1021/acscatal.3c05302.

(57) Stewart, N. K.; Toth, M.; Quan, P.; Beer, M.; Buynak, J. D.; Smith, C. A.; Vakulenko, S. B. Restricted Rotational Flexibility of the C5α-Methyl-Substituted Carbapenem NA-1-157 Leads to Potent Inhibition of the GES-5 Carbapenemase. ACS Infectious Diseases 2024, 10 (4), 1232–1249. DOI: 10.1021/acsinfecdis.3c00683.

(58) Twidale, R. M.; Hinchliffe, P.; Spencer, J.; Mulholland, A. J. Crystallography and QM/MM Simulations Identify Preferential Binding of Hydrolyzed Carbapenem and Penem Antibiotics to the L1 Metallo-β-Lactamase in the Imine Form. Journal of Chemical Information and Modeling 2021, 61 (12), 5988–5999. DOI: 10.1021/acs.jcim.1c00663. Borbulevych, O. Y.; Martin, R. I.; Westerhoff, L. M. The critical role of QM/MM X-ray refinement and accurate tautomer/protomer determination in structure-based drug design. Journal of Computer-Aided Molecular Design 2021, 35 (4), 433–451. DOI: 10.1007/s10822-020-00354-6.

(59) Smith, C. A.; Nossoni, Z.; Toth, M.; Stewart, N. K.; Frase, H.; Vakulenko, S. B. Role of the Conserved Disulfide Bridge in Class A Carbapenemases. J Biol Chem 2016, 291 (42), 22196–22206. DOI: 10.1074/jbc.M116.749648 From Nlm.

(60) Le Terrier, C.; Nordmann, P.; Freret, C.; Seigneur, M.; Poirel, L. Impact of Acquired Broad Spectrum β-Lactamases on Susceptibility to Novel Combinations Made of β-Lactams (Aztreonam, Cefepime, Meropenem, and Imipenem) and Novel β-Lactamase Inhibitors in Escherichia coli and Pseudomonas aeruginosa. Antimicrob Agents Chemother 2023, 67 (7), e0033923. DOI: 10.1128/aac.00339-23 From NLM.

