## Supplementary Information for "Mechanistic Basis for Inhibition of the Extended Spectrum Class A β-Lactamase GES-1 by Tazobactam and Enmetazobactam"

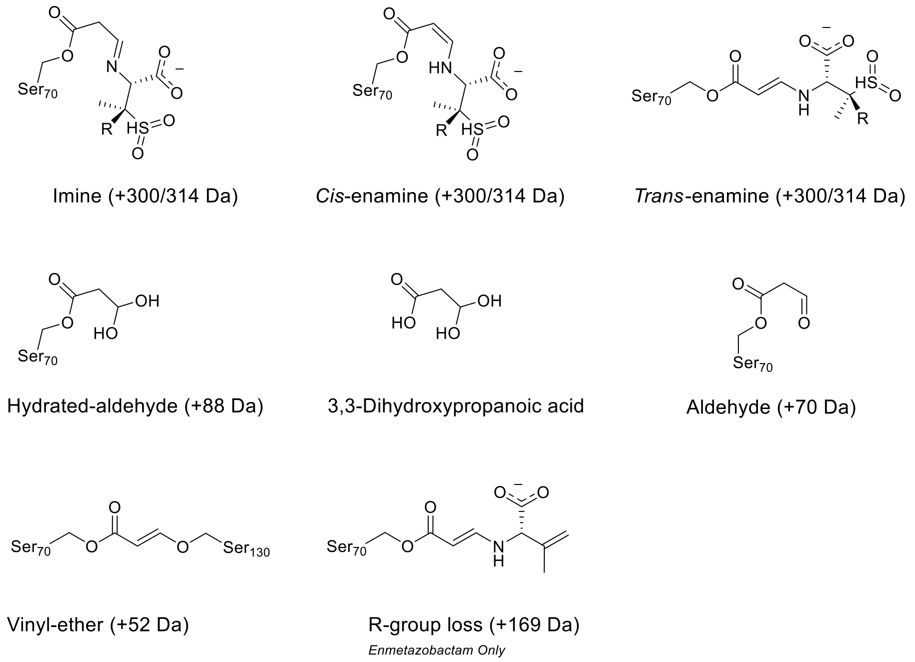


***Figure S1: Breakdown Products of PAS Inhibitors After Exposure to Class A β-Lactamases.*** *R denotes attachment of substituents differentiating sulbactam, tazobactam and enmetazobactam. The expected mass increase that would be observed by mass spectrometry experiments if each product was acylated to the enzyme is shown in brackets. 3,3-Dihydroxypropanoic acid is the (non-covalent) deacylation product of the hydrated aldehyde, and consequently is not associated with any mass increase compared to uncomplexed enzyme.*

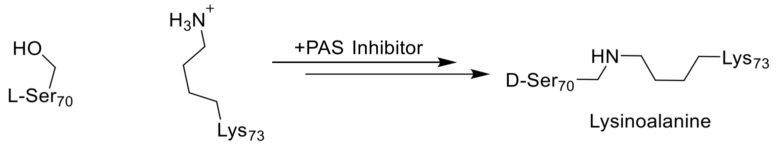


***Figure S2: Formation of Active-Site Lysinoalanine Cross-Link.*** *The mechanism of enzyme-catalysed formation of lysinoalanine from the Ser70-(enme)tazobactam acyl-enzyme complex is unknown, but is thought to include L- to D- epimerisation of Ser70 and possible dehydroalanine formation (1, 2).*

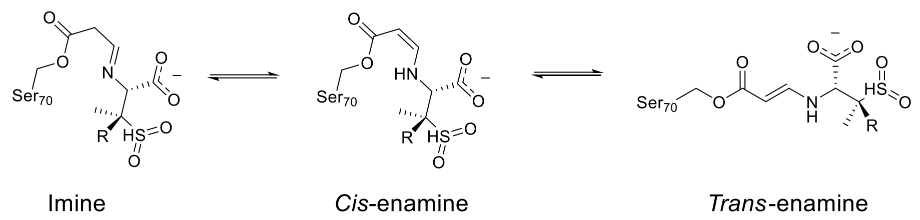


***Figure S3: Tautomerisation of PAS Inhibitor Acyl-Enzyme Complexes.*** *R denotes attachment of substituents differentiating sulbactam, tazobactam and enmetazobactam.*

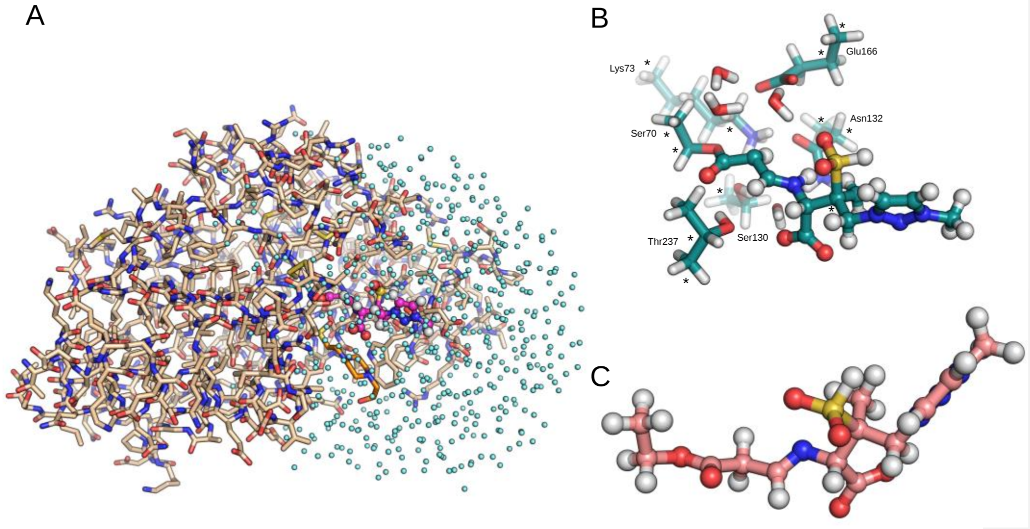


***Figure S4:QM/MM Set Up.*** *Representative structures for the QM/MM (A), active site model (B) and acyl-adduct model (C) geometry optimisations. Water molecules in the QM/MM starting structure are shown as cyan spheres. In (A) atoms treated by QM (Ser70 Cβ, Ser70 Oγ and enmetazobactam-derived covalent adduct) are shown in pink, whilst bound HEPES is shown in orange. For clarity, all hydrogen atoms other than those present in the non-covalent adduct are removed. In (B) asterisks denote atoms that were held fixed during geometry optimisation, to preserve crystallographic geometry.*

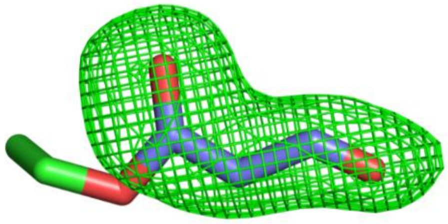

***Figure S5: Omit Density for GES-1 Bound Tazobactam.*** *$F_o$-$F_c$ omit map calculated after removal of ligand is shown contoured at 3σ around atoms of the tazobactam-derived covalent adduct only. Tazobactam carbon atoms are shown blue, GES-1 (Ser70) carbon atoms in green.*

**Commented [JS1]:** Correct?

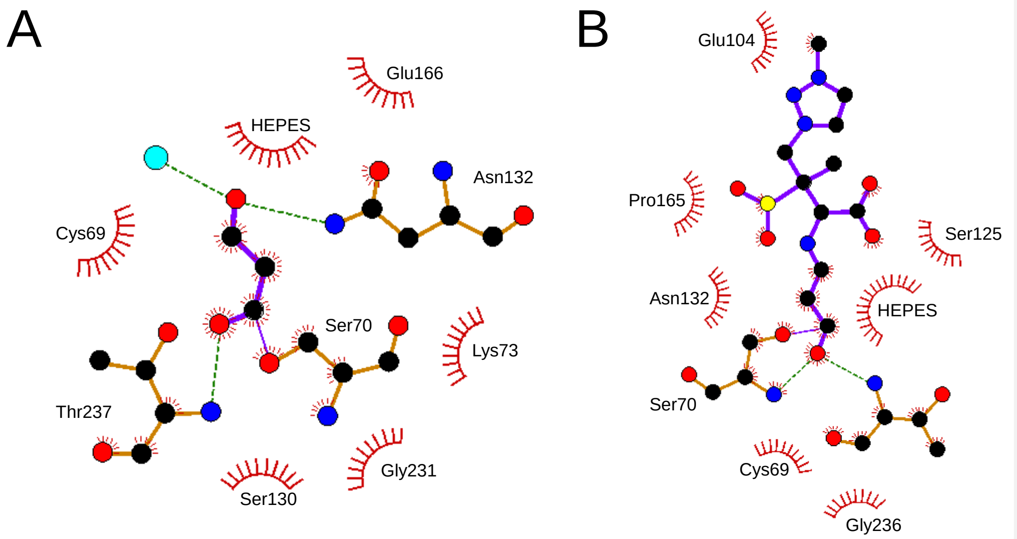


***Figure S6: Interactions Between GES-1 and PAS Inhibitor-Derived Covalent Adducts.*** *Interaction diagrams of tazobactam- (A) and enmetazobactam- (B) derived GES-1 acyl-enzymes. Hydrogen bonds are displayed as green dashed lines, and water molecules as cyan spheres.*

**Commented [JS2]:** Citation if this was LigPlot-derived, or did you draw yourself?

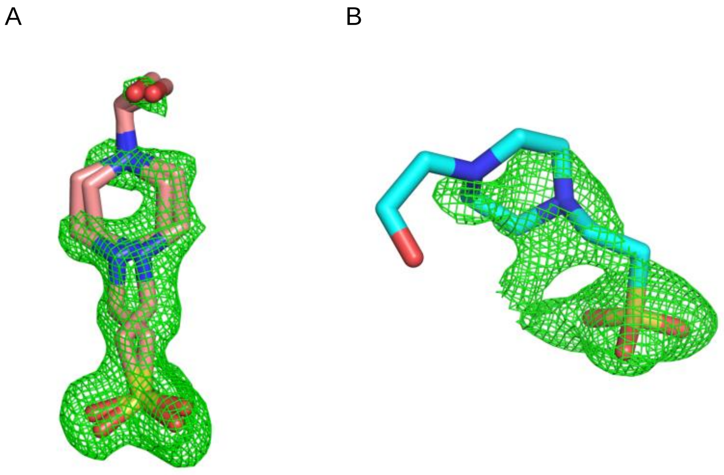


***Figure S7: Omit Density for Bound HEPES.*** *$F_o$-$F_c$ omit maps (green mesh) calculated after removal of ligand and contoured at 3σ around HEPES atoms in the enmetazobactam-derived (A) and tazobactam-derived (B) acyl-enzymes. Representative images shown here are taken from the active sites of chain A of both structures.*

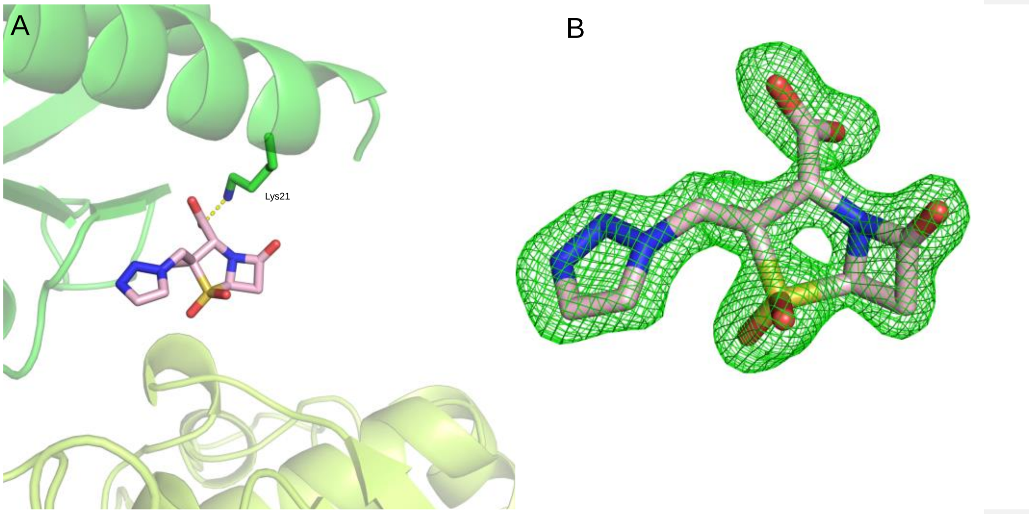


***Figure S9:Binding of Intact Tazobactam at Crystallographic Interface.*** *A) Orientation of tazobactam bound at a crystal symmetry interface (lime green chain (bottom) represents the symmetry partner of the green chain (top)). The tazobactam carboxylate moiety is within hydrogen-bonding distance of the Lys21 side chain amide. B)* $F_o$*-*$F_c$ *omit map (green mesh, calculated after removal of ligand) around bound tazobactam, contoured at 3σ.*

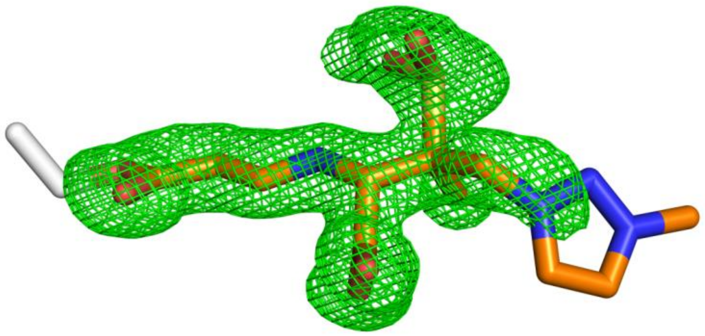

***Figure S10: Omit Density for Covalently Bound Enmetazobactam-Derived Molecule.*** *$F_o$-$F_c$ omit map (green mesh, calculated after removal of ligand) contoured at 3σ from chain A. Density is similar in chain B.*

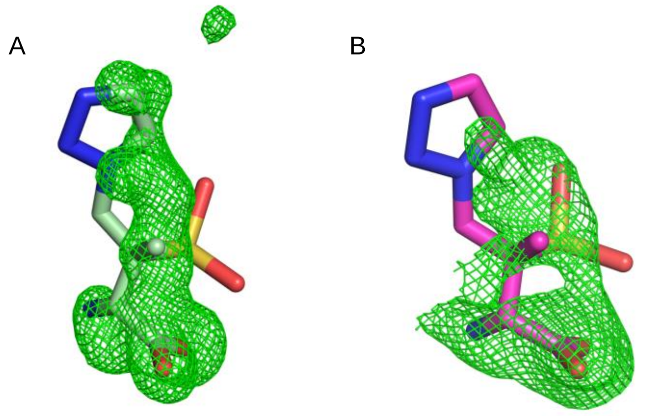


***Figure S11: Omit Density for Bound HEPES in Comparison with Previously Modelled GES-2:Tazobactam Breakdown Product.*** *$F_o$-$F_c$ omit map (green mesh), calculated after removal of ligand and contoured around bound HEPES in chains A of enmetazobactam-derived (A) and tazobactam-derived(B) acyl-enzymes; superposed upon previously described tazobactam-derived breakdown product observed in crystal structure of a GES-2 complex (PDB 3NIA (3)). Tazobactam-derived product was aligned to each structure separately. Alignments strongly indicate that the previously modelled compound, identified within the active site of GES-2 (GES-1 G170N point variant) after exposure to tazobactam is not a good fit to the electron density in either of the structures reported here. In both cases the $F_o$-$F_c$ omit map density is similar in chain B.*

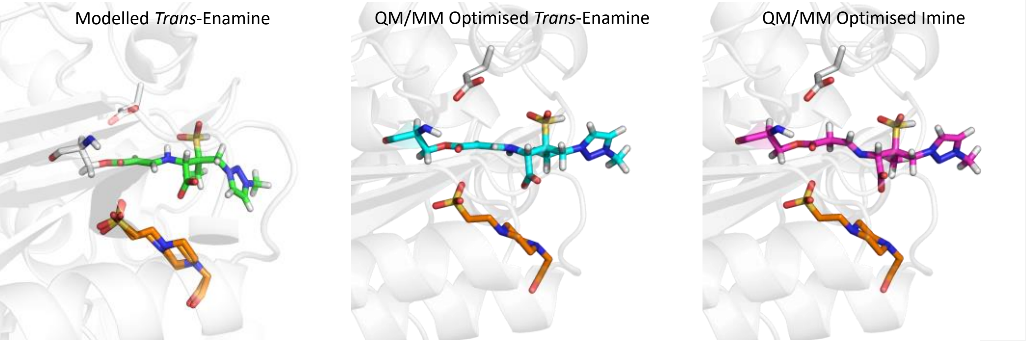


***Figure S12: Models of Enmetazobactam-Derived Species in GES-1 Complex Structures.*** *A) Crystallographic model of trans-enamine; B) QM/MM-optimised trans-enamine; C) QM/MM-optimised imine. Optimised trans-enamine (B), centre) more closely resembles the structure modelled into the experimental electron density (left) than does the imine (right). Note that the methyltriazole ring (right) is poorly defined by experimental electron density, consistent with the movement of this moiety during QM/MM geometry optimisation.*

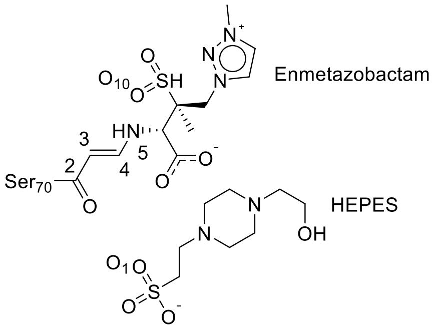


***Figure S13: Atom Numbering of Enmetazobactam and HEPES.*** *Atom numbers shown are those used in the angle and distance analysis table (Table S2).*

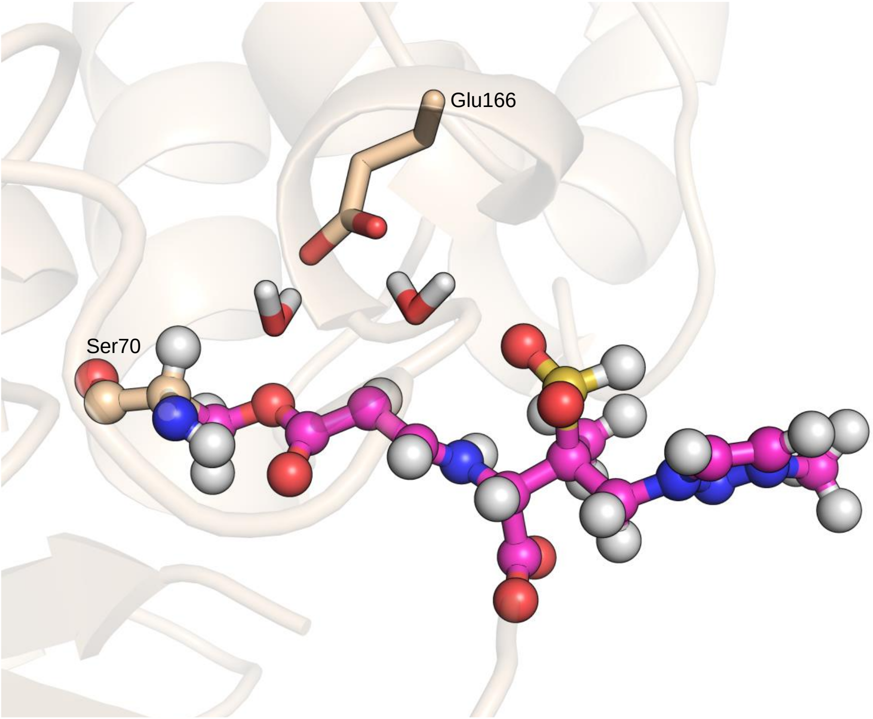


***Figure S14**: **Proximal Water Molecules Affect Relative Stability of the Trans-Enamine.** QM region (enmetazobactam-derived acylated compound and Ser70 Oγ and Cβ, including bound hydrogens) is shown in magenta. Two water molecules (sticks) affect the relative stability of the trans-enamine, depending on hydrogen-bonding interactions with the Glu166 side chain and the sulphone moiety of the enmetazobactam-derived acylated compound. The lowest energy orientation (calculated using both the QM and MM regions), compared to structures starting with different orientations of the two water molecules, is shown. QM/MM geometry optimisation were completed using the B3LYP/6-31G (d) level of theory with Grimme's D3 dispersion correction and Becke-Johnson damping in the QM region. B3LYP-D3(BJ)/def2-TZVP was used in the QM region for single point energy calculations of the optimised structures.*

*Table S1: Crystallographic Statistics*

| | GES-1:enmetazobactam | GES-1:tazobactam | Uncomplexed GES-1 (enmetazobactam soaking condition) | Uncomplexed GES-1 (tazobactam soaking condition) |
|---|---|---|---|---|
| **PDB Code** | **9ENX** | **9ENY** | **9ENW** | **9ENV** |
| **Data Collection** | | | | |
| Wavelength (Å) | 0.73379 | 0.97628 | 0.81530 | 0.7838 |
| Resolution Range | 81.04 – 1.23 (1.25-1.23) | 63.90 – 1.30 (1.32-1.30) | 52.71-1.66 (1.69-1.66) | 43.89-1.60 (1.63-1.69) |
| Space Group | *P* 21 | *P* 21 21 21 | *P* 21 | *P* 21 21 21 |
| Molecules/ASU | 2 | 2 | 2 | 2 |
| Cell Dimensions | | | | |
| a, b, c (Å) | 42.80, 81.03, 71.55 | 75.95, 80.71, 104.55 | 42.63, 80.610, 71.07 | 75.56, 80.81, 104.54 |
| α, β, γ (°) | 90, 90, 90 | 90, 90, 90 | 90, 90, 90 | 90, 90, 90 |
| Multiplicity | 6.9 (6.8) | 13.2 (13.4) | 6.8 (6.9) | 14.0 (13.9) |
| Completeness (%) | 99.8 (96.0) | 100.0 (100.0) | 100.0 (100.0) | 100.0 (99.0) |
| I/σ(I) | 8.1 (0.3) | 12.6 (0.6) | 5.4 (0.7) | 6.9 (0.4) |
| $R_{pim}$ | 0.045 (1.151) | 0.026 (1.331) | 0.085 (1.151) | 0.064 (1.083) |
| $CC_{1/2}$ | 0.997 (0.314) | 0.999 (0.340) | 0.995 (0.249) | 0.997 (0.324) |
| **Refinement** | | | | |
| Resolution | 70.04-1.23 | 61.42-1.36 | 41.79-1.66 | 43.89-1.60 |
| No. reflections | 135565 | 115448 | 55561 | 84864 |
| R-work/R-free | 0.1519/0.1935 | 0.1516/0.1946 | 0.2022/0.2511 | 0.1777/0.2113 |
| No. non-H atoms | | | | |
| Protein | 4096 | 4102 | 4044 | 4126 |
| Solvent | 674 | 644 | 520 | 626 |
| Ligand | 72 | 30 | - | - |
| Average B-Factors | | | | |
| Protein | 19.8 | 30.78 | 25.7 | 27.5 |
| Solvent | 35.2 | 35.6 | 32.7 | 37.9 |
| Ligand | 27.9 | 33.7 | - | - |
| R.m.s Deviations | | | | |
| Bond Lengths (Å) | 0.008 | 0.008 | 0.01 | 0.01 |
| Bond Angles (°) | 0.959 | 0.982 | 1.052 | 0.988 |
| Ramachandran (%) | | | | |
| Outliers | 0.0 | 0.0 | 0.0 | 1.6 |
| Favoured | 98.1 | 98.9 | 97.5 | 98.1 |

***Table S2: Comparison of Distances and Bond Angles in Models of the Enmetazobactam-Derived Species.*** *Measurements are shown for crystallographic model of bound ligand and QM-optimised trans-enamine and imine species. (Note that HEPES was not included in the cluster or small QM models, so no values are reported for the HEPES$^{O1}$-N5 distance.) See Figure S13 for atom numbering.*

| **Analysis** | **X-ray Modelled *Trans*-Enamine** | **X-ray Modelled Imine** | **QM/MM Optimised *Trans*-Enamine** | **QM/MM Optimised Imine** | **Cluster Model *Trans*-Enamine** | **Cluster Model Imine** | **Acyl-Adduct Model *Trans*-Enamine** | **Acyl-Adduct Model Imine** |
|---|---|---|---|---|---|---|---|---|
| C2-C3-C4 Angle (°) | 120.9 | 119.5 | 120.6 | 114.4 | 121.3 | 116.2 | 119.2 | 112.1 |
| C3-C4-N5 Angle (°) | 125.2 | 120.0 | 124.4 | 121.1 | 123.9 | 120.7 | 126.1 | 120.7 |
| C2-C3-C4-N5 Torsion Angle(°) | −172.8 | −178.2 | 177.3 | 125.8 | 173.3 | 149.3 | −178.9 | −125.7 |
| HEPES$^{O1}$-N5 Distance (Å) | 4.7 | 4.6 | 4.6 | 4.1 | - | - | - | - |
| O10-N5 Distance (Å) | 2.7 | 2.7 | 2.9 | 3.1 | 2.9 | 2.9 | 3.0 | 2.9 |

***Table S3: Relative Energies of Tautomeric Species from QM and QM/MM Calculations.*** *Values for modelled imines were used as reference energies. In each case the level of theory for the geometry optimisation was B3LYP 6-31G (d) with Grimme's D3 dispersion correction and Becke-Johnson damping whilst B3LYP-D3(BJ)/def2-TZVP was used for single point energy calculations.*

| | **Relative Energy (kcal mol$^{-1}$)** | | |
|---|---|---|---|
| **Tautomer State** | **QM/MM** | **Active Site Model** | **Acyl-Adduct Only** |
| Imine | 0 | 0 | 0 |
| *Trans*-Enamine | –3.3 | –9.2 | –5.7 |

### Supplementary References


1. Hinchliffe P, Tooke CL, Bethel CR, Wang B, Arthur C, Heesom KJ, et al. Penicillanic Acid Sulfones Inactivate the Extended-Spectrum β-Lactamase CTX-M-15 through Formation of a Serine-Lysine Cross-Link: an Alternative Mechanism of β-Lactamase Inhibition. mBio. 2022;13(3):e01793-21.
2. Lang PA, Raj R, Tumber A, Lohans CT, Rabe P, Robinson CV, et al. Studies on enmetazobactam clarify mechanisms of widely used β-lactamase inhibitors. Proceedings of the National Academy of Sciences. 2022;119(18):e2117310119.
3. Frase H, Smith CA, Toth M, Champion MM, Mobashery S, Vakulenko SB. Identification of products of inhibition of GES-2 beta-lactamase by tazobactam by x-ray crystallography and spectrometry. J Biol Chem. 2011;286(16):14396-409.
